# Whole Genome Sequencing Reveals Autooctoploidy in the Chinese Sturgeon and its Evolutionary Trajectories

**DOI:** 10.1101/2023.06.06.543648

**Authors:** Binzhong Wang, Bin Wu, Xueqing Liu, Yacheng Hu, Yao Ming, Mingzhou Bai, Juanjuan Liu, Kan Xiao, Qingkai Zeng, Jing Yang, Hongqi Wang, Baifu Guo, Chun Tan, Zixuan Hu, Xun Zhao, Yanhong Li, Zhen Yue, Junpu Mei, Wei Jiang, Yuanjin Yang, Zhiyuan Li, Yong Gao, Lei Chen, Jianbo Jian, Hejun Du

**Affiliations:** Hubei Key Laboratory of Three Gorges Project for Conservation of Fishes, Yichang, Hubei, 443100, China; Chinese Sturgeon Research Institute, China Three Gorges Corporation, Yichang, Hubei, 443100, China; Yangtze River Biodiversity Research Center, China Three Gorges Corporation, Wuhan, Hubei, 430014, China; BGI-Shenzhen, Shenzhen, 518083, China; BGI Genomics, BGI-Shenzhen, Shenzhen, 518083, China; BGI-Sanya, BGI-Shenzhen, Sanya, 572025, China; Department of Biotechnology and Biomedicine, Technical University of Denmark, Lyngby, 2800, Denmark

**Keywords:** Chinese sturgeon, Whole-genome sequencing, Autooctoploid, Polyploidization and diploidization, Whole genome duplication

## Abstract

The Order Acipenseriformes, which include sturgeons and paddlefishes, represent “living fossils” with complex genomes that are good models for understanding whole genome duplication (WGD) and ploidy evolution in fishes. Here we sequenced and assembled the first high-quality chromosome-level genome for the complex octoploid *Acipenser sinensis* (Chinese sturgeon), a critically endangered species that also represents a poorly understood ploidy group in Acipenseriformes. Our results show that *A. sinensis* is a complex autooctoploid species containing four kinds of octovalents (8 n), a hexavalent (6 n), two tetravalents (4 n), and a divalent (2 n). We propose based on an analysis taking into account delayed rediploidization that its octoploid genome composition results from two rounds of homologous whole genome duplications (WGDs), and further provide insight into the timing of its ploidy evolution. This study provides the first octoploid genome resource of Acipenseriformes for understanding ploidy composition and evolutionary trajectories of polyploidy fishes.

## Introduction

The order Acipenseriformes, which include sturgeon and paddlefishes, is an ancient group of fishes with wide distribution in the Northern Hemisphere. Many Acipenseriformes are threatened or endangered, particularly due to their commercial value for meat and caviar. As “living fossils”, Acipenseriformes retain primitive characteristics (such as a heterocercal tail and cartilaginous skeleton) and occupy the basal position of Actinopterygii phylogeny [1]. They also exhibit a slow evolution rate [2], complex genomes (half of their chromosomes are micro-chromosomes), and complex ploidy, with the order divisible into three ploidy classifications: Group A: ∼120 chromosomes, nuclear DNA content of 3.2–4.6 pg, Group B: ∼240 chromosomes, nuclear DNA content of 6.1–9.6 pg, and Group C: ∼360 chromosomes, nuclear DNA content of 13.1–14.2 pg [3, 4]. A better understanding of Acipenseriformes could aid in conservation efforts and provide insights into the understanding of whole genome duplication (WGD) and ploidy evolution in fishes [5-7].

WGDs are very common in the evolution of fish [8-10] and subsequent rediploidization further increases the complexity of the genomes. Both Acipenseriformes and teleosts (ray-finned fishes except primitive bichirs, sturgeons, paddlefishes, freshwater garfishes, and bowfins) have undergone at least three rounds of WGDs. The first two rounds include the first round WGD (1R) that occurred at ∼600 million years ago (MYA) and a jawed vertebrate-specific second round WGD (2R) that occurred after the divergence of lamprey and jawed vertebrates. Teleosts then underwent a teleost-specific third round (Ts3R) [11-14], whereas Acipenseriformes underwent an independent Acipenseriforme-specific WGD (As3R) [11]. Acipenseriformes are thought to have undergone a delayed rediploidization, in which a species radiates an extensive time after a WGD (i.e. a timescale on the order of millions of years) [15], resulting in complex ploidies. However, the ploidy compositions of most Acipenseriformes species have been challenging to clarify. The ploidy debates, for example whether Groups A and B are diploid and tetraploid [5] or are instead tetraploid[16] and octoploid [17-19], or even are paleotetraploidy versus modern/functional diploidy in the case of Group A [4], have lasted for half a century [20, 21]. It has been difficult to end the debate solely by relying on traditional DNA content measurement, cytogenetics, and molecular biology techniques [18, 22-24], and thus whole-genome sequencing analyses is needed to help resolve outstanding question. Moreover, efforts in estimation of polyploidization and speciation time need to appropriately take into account rediploidization effects.

The ‘Lineage-specific Ohnologue Resolution’ (LORe) model (Figure S1A) was proposed to address delayed rediploidization of sister lineages that share the common WGD [15]. In the LORe model, speciation precedes rediploidization, allowing for independent ohnologue divergence in sister lineages that shared an ancestral WGD event. A phylogenetic implication of LORe is the absence of 1:1 orthology between ohnologue pairs from different lineages, leading to the definition of the term ‘tetralog’ to describe a 2:2 homology relationship between ohnologues in sister lineages. Order differences in divergency and polypolidization, as well as the influence of species characteristics, will lead to differential enrichment of LORe and ‘Ancestral Ohnologue Resolution’ (AORe) [15, 25]. Thus, the estimation of polyploidization and speciation time that ignore the coefficient of LORe and AORe would produce an inaccurate result based on the traditional methods of globally homologous comparison. Previous reports on WGD and rediploidization processes only focused on Acipenseriformes species of group A but not on species of Group B mainly due to a lack of Group B genomic resources and the influence of delayed rediploidization on the analyses was not considered [11, 26], although the phylogenetic analysis of autopolyploid American paddlefish genome studies based on traditional methods of globally homologous comparison did suggest an influence of delayed rediploidization [26].

To address the lack of Group B resources and to further clarify ploidy in Acipenseriformes species, we examined the Group B Acipenseriformes species *Acipenser sinensis* (Chinese sturgeon) by whole genome sequencing. *A. sinesis* is a critically endangered large fish in China and also a sturgeon species of world-wide concerned with the lowest distribution latitude [27]. *A. sinesis* has ∼264 chromosomes, including 124 macro-chromosomes and ∼140 micro-chromosomes [18, 28], and is also considered a paleooctoploid [29]. From our whole genome sequencing and subsequent analysis, we now report the first high-quality genome assembly of *A. sinensis* and its ploidy compositions of post-rediploidization. Furthermore, by combining *A. sinensis* (Acipenseridae, group B), *Acipenser ruthenus* (Acipenseridae, group A) [30], and *Polyodon spathula* (Polyodontidae, group A) [26] genomic data, and integrating assessment of the influence of the delayed rediploidization, we also uncover the evolutionary trajectories of *A. sinensis*.

## Results

### *A. sinensis* genome sequencing, assembly, and annotation

To obtain a high-quality genome assembly, DNA from a meiotic gynogenetic male *A. sinensis* was sequenced by combining the Illumina platform, the PacBio platform, and chromosome conformation capture (Hi-C) sequencing technology. We obtained 421.58 Gb clean Illumina short-read data (Table S1), 221.96 Gb clean PacBio long-reads data (Table S2), and 172.87 Gb clean Hi-C data (Table S3). Illumina and PacBio reads were assembled into the initial contigs of ∼1.99 Gb with an N50 size of ∼4.07 Mb (**Table 1**). Clean Hi-C reads were applied to anchor contigs into 66 scaffolds corresponding to 66 chromosomes of two monoploid genomes (**Figure 1**A and B; **Table 1**). The final genome assembly was 1.99 Gb with a scaffold N50 size of ∼48.46 Mb, and 98.3% of assembled sequences were assigned to chromosomes (**Figure 1**C; **Table 1**). This genome assembly size was comparable to the size estimated by a *k*-mer-based method (1.975 Gb) and a quarter of the DNA content (2.27 pg/2C) estimated by the Flow Cytometer (Figures S2 and S3; Table S4). We observed a high correlation (R^2^ = 0.98) between 66 assembled chromosome sizes and the relative physical length of chromosomes based on our karyotype results (**Figure 1**D; Table S5). The completeness of genome assembly was 95.6%, including 60.0% of the ‘complete and single-copy BUSCOs’ and 35.6% of the ‘complete and duplicated BUSCOs’ (Table S6). We further determined that the mapping rate and coverage of the sequences are 99.67% and 93.62%, respectively, using the BWA alignment results with 500 bp data.

**Figure 1.**
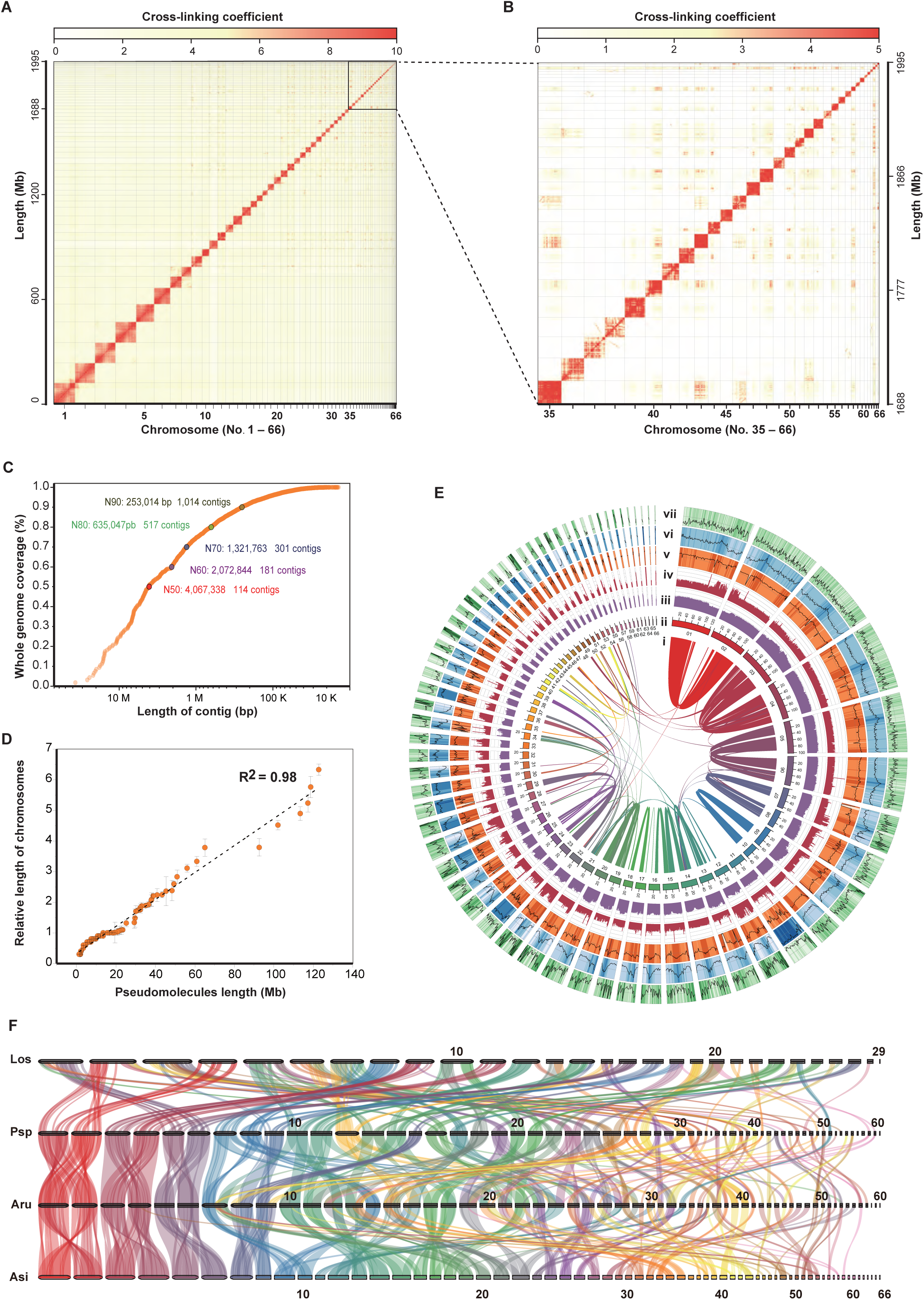
Genome assembly and evaluation of *Acipenser sinensis*. **A.** Heatmap of interactions within and among chromosomes based on Hi-C data. Each dark red block with a high cross-linking coefficient and clear boundaries is a chromosome region. Sixty-six chromosomes were segmented. **B**. Mosaics of 33 detailed micro-chromosomes (35–66) were displayed. **C.** Length distribution of contigs in the whole-genome assembly. The key indicators: N50 and the number of contigs are 4,067,338 bp and 114, respectively. N90 and the number of contigs are 253,014 bp and 1,014, respectively. **D.** Correlation analysis between pseudomolecule lengths and relative length of chromosomes (R^2^ = 0.98, *P* = 2.97E-49). **E**. Features of 66 assembled chromosomes. Tracks from the inner to the outside are indicated as follows: I, the relationship between collinearity blocks; II, pseudo-chromosomes; III, frequencies of suballele (0–0.5); IV, sequencing depths (0–200×); V, GC contents (20%–50%); VI, TE, transposable element contents (0–100%); and VII, gene contents (0-50). **F**. Relationships of collinearity blocks among *Lepisosteus osseus* (Los), *Polyodon spathula* (Psp), *Acipenser ruthenus* (Aru), and *Acipenser sinensis* (Asi).

**Table 1.** Statistics of genome assembly of three sequenced Acipenseriforme species.

| Parameter | *Acipenser sinensis* | *Acipenser ruthenus* [1] | *Polyodon spathula* [2] |
| --- | --- | --- | --- |
| Ploidy | 8 | 4 | 4 |
| Assembly size (bp) | 1,995,374,126 | 1,830,501,248 | 1,542,083,420 |
| Contig N50 (bp) | 4,067,338 | 600,839 | 3,441,286 |
| Scaffold N50 (bp) | 48,462,029 | 42,361,471 | 48,906,729 |
| Contig N90 (bp) | 253,014 | 72,599 | 255,888 |
| Scaffold N90 (bp) | 13,550,145 | 3,341,419 | 7,546,224 |
| Length of gaps (bp) | 1,560,502 | 1,254,212 | 1,696,466 |
| GC content | 40.76% | 39.76% | 39.02% |
| No. of chromosomes | 66 | 60 | 60 |
| Chromosome length (bp) | 1,962,255,781 | 1,651,776,498 | 1,487,634,427 |
| Chromosome anchoring rate | 98.3% | 90.2% | 96.5% |
| Complete BUSCOs of assembly | 95.60% | 98.30% | 93.7% |
| No. of coding genes number | 36,837 | 35,379 | 30,260 |
| Complete BUSCOs of genes | 93.10% | - | - |

We integrated *de novo*, protein homology-based, and RNA-seq data-assisted methods to predict gene structures (Table S7). A total of 36,837 protein-encoding genes were predicted with an average length of 14.3 Kb (**Table 2**). BUSCO evaluation indicated that the annotation covered 93.1% of the vertebrate core gene sets (Table S8). We annotated 34,950 protein-coding genes (94.88% of the predicted genes), and 91.78%, 82.36%, 78.18%, 69.09%, 91.18%, 84.21%, and 57.81% genes matched to Non-Redundant Protein Sequence Database in NCBI (Nr), Swissprot, KEGG, KOG, TrEMBL, Interpro, and GO databases, respectively (Figure S4; Table S9). Meanwhile, we identified 991.35 Mb repetitive elements, which accounted for 49.68% of the *A. sinensis* genome, containing 204.28 Mb tandem repeats (10.23% of the genome) (Table S10) and 903.50 Mb transposable elements (TEs) (45.28% of the genome). Among the assembled genome, 16.34% are long interspersed nuclear elements (LINEs), 17.43% are long terminal repeats (LTRs), and 1.88% are short interspersed nuclear elements (SINEs), and 17.67% are class II TEs (DNA transposons) (Table S11). Overall, we have sequenced and assembled a high-quality genome of *A. sinensis*.

**Table 2.** General statistics of predicted protein-coding genes.

| Type | Gene set | Number | Ave gene length (bp) | Ave CDS length (bp) | Ave exon count per gene | Ave exon length (bp) | Ave intron length (bp) |
| --- | --- | --- | --- | --- | --- | --- | --- |
| *De novo* | Augustus | 105,618 | 10,958.13 | 1,231.69 | 4.93 | 249.99 | 2,476.87 |
|  | GeneScan | 132,324 | 14,471.77 | 1,495.38 | 5.95 | 251.45 | 2,623.12 |
| Protein-homology | *Callorhinchus milii* | 88,902 | 6,333.78 | 923.12 | 3.7 | 249.72 | 2,006.53 |
|  | *Danio rerio* | 94,544 | 6,991.85 | 1,051.28 | 4.06 | 259.05 | 1,942.49 |
|  | *Latimeria chalumnae* | 140,860 | 4,618.43 | 876.97 | 3.13 | 280.21 | 1,756.83 |
|  | *Lepisosteus oculatus* | 102,019 | 7,265.95 | 1,007.46 | 4.15 | 242.63 | 1,985.44 |
|  | *Petromyzon marinus* | 94,541 | 3,151.53 | 762.17 | 2.37 | 322.15 | 1,749.33 |
| RNA | RNA | 23,727 | 6,467.33 | 1,097.87 | 6.42 | 170.93 | 1,092.26 |
|  | Gene set | 36,837 | 14,306.64 | 1,392.56 | 8.03 | 173.37 | 1,708.37 |

*Note:* The average transcript length did not contain UTR. Three approaches, Homolog (*Callorhinchus milii*, *Danio* *rerio*, *Latimeria chalumnae*, *Lepisosteus oculatus*, *Petromyzon marinus*), *De novo* (GenScan, AUGUSTUS), and RNA, were employed in gene prediction. Results of Homolog and *De novo* were consolidated using the program GLEAN.

### Collinearity analysis and phylogenetic tree construction

To analyze the genome assembly quality and chromosome evolution, we performed a collinearity analysis among the genomes of three sequenced Acipenseriforme species and a close relative species of Acipenseriformes on the evolutionary tree, including *L. oculatus*, *P. spathula*, *A. ruthenus*, and *A. sinensis*. A total of 25,687 collinear genes were detected in the *A. sinensis* internal genome. We also obtained collinear genes between *A. sinensis* and *A. ruthenus* (31,348 genes), *A. sinensis* and *P. spathula* (28,945 genes), *A. sinensis* and *L. oculatus* (16,062 genes), *A. ruthenus* and *P. spathula* (34,031 genes), and *P. spathula* and *L. oculatus* (18,053 genes) (Table S12). The six relatively large chromosomes displayed the definite two-to-two collinearity relationship between *A. sinensis*, *A. ruthenus*, and *P. spathula* (**Figure 1**F).

Meanwhile, 21,410 gene families were obtained by clustering homologous gene sequences among 13 species. We chose these fish species for phylogenetic analysis based on three principles: (1) that they have sequenced whole genomes, (2) that they represent important branches or nodes within the phylogenetic tree, and (3) that they are typical representative species or model species. According to these three principles, *Petromyzon marinus* represents jawless fishes, *Callorhinchus milii* represents cartilaginous fishes, *Latimeria chalumnae* represents ancient fishes and lobe-finned fishes, *Polypterus senegalus* represents polypterids, *Lepisosteus oculatus* represents the evolution node species of Actinopterygian and bony fishes, *Salmon salar* represents a highly tetraploid species, and *Cyprinus carpio* represents typical allopolyploid fishes, *Gadus morhua* represents typical bony fish, and *Polyodon spathula* represents paddlefish. *A. ruthenus* represents the group A population of sturgeon in the Atlantic branch of the sturgeon family (with a chromosome number of ∼120) and *A. sinensis* represents the Group B species of sturgeon in the Pacific branch of the sturgeon family (with a chromosome number of ∼240). Lastly, *Oryzias latipes* and *Danio rerio* were included as important model organism species.

We constructed the phylogenetic tree with 2,096 genes using two methods, PhyML [31] (Table S13) and Astral [32] (Figure S5), which both yielded a phylogenetic tree with the same topological structure. The resulting trees revealed that *A. sinensis* and *A. ruthenus* have the same Acipenseridae ancestor, whereas *P. spathula* is a sister lineage with the Acipenseridae and classified as Polyodontidae. The trees also showed that as a cluster of representative ancient species, the Acipenseriformes and Lepisosteiformes diverged from the same evolutionary branch (Figure S6). The results are consistent with the previous studies [33] and thus together with the collinearity analysis, verify the high assembly quality and integrity of our genomic dataset and analysis.

### Ploidy identification

Traditional karyotype analyses have shown that *A. sinensis* has 264 chromosomes (Figure S7), approximately four times the 60-chromosome karyotype of the common diploid ancestor of Acipenseriformes [18, 28, 34]. Thus, *A. sinensis* was presumed to be an octoploid species. To further explore *A. sinensis* ploidy at the genome level, we obtained 200 Gb genome sequences of a normal reproductive animal, instead of a meiotic gynogenetic animal, using the BGISEQ platform. We analyzed genome-wide simple sequence repeats (SSRs) and single nucleotide polymorphisms (SNPs) based on the data. SSR analyses showed that the largest number of alleles at a single locus was up to eight (Figure S8; Table S14), implying that the species has eight homologous chromosomes.

We further identified SNPs in *A. sinensis* (22,324,005 SNPs) and *A. ruthenus* (9,826,321 SNPs) for comparative analysis of ploidy. Heterozygosity results indicated that *A. sinensis* (1.12%) is approximately two folds of tetraploidy *A. ruthenus* (0.54%) (Table S15). We evaluated and plotted the allele frequency and ploidy (n) of the SNP sites. In *A. ruthenus*, most of the SNP allele frequencies displayed 1/2 and 1/4, but both depths pointed to 4 n (∼46×) (**Figure 2**A), which indicates that the genotypes were AABB and AAAB. This implies that *A. ruthenus* experienced an autotetraploidization event followed by a rediploidization event, which is similar to the conclusions from a previous study by Du et al. [11]. *A. sinensis* has more complex ploidy, and so bivalents (2 n), tetravalents (4 n), and octovalents (8 n) were all observed in the analysis (**Figure 2**B). *A. sinensis* exhibited four peaks in the SNP frequency curve, whereas *A. ruthenus*, the tetraploid sturgeon, only exhibited two peaks. The main peak of ploidy in *A. sinensis* was 4 n, pointing to 1/4 and 1/2 allele frequency. Most importantly, we detected the first peak at the position of 1/8 in *A. sinensis* allele frequency. This peak mostly pointed to 8n ploidy, suggesting that the eight monoploids have high similarity and revealed the octoploid features of *A. sinensis*.

**Figure 2.**
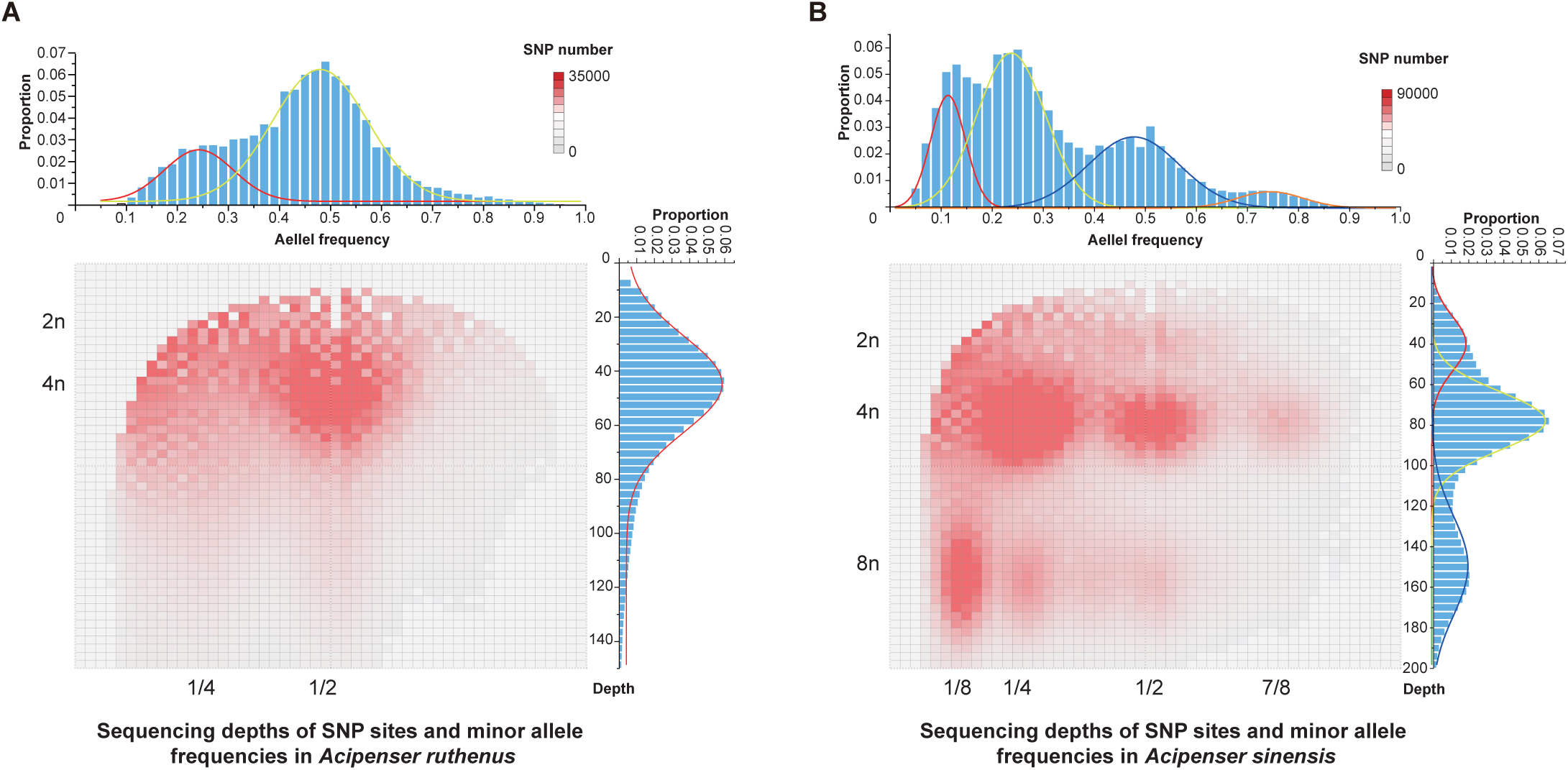
**Sequencing depths of SNP sites and minor allele frequencies in the assemblies of *Acipenser sinensis* and *Acipenser ruthenus***. In the main plot, the X and Y axes represent minor allele frequency and SNP sequencing depth, respectively. The right and upper bar charts show the number of hits at the given frequency and depth. Multimodal Gaussian fitting shows: **A.** *A. ruthenus* has two significant peaks of the minor allele frequency at 1/4 and 1/2, respectively, and SNP site sequencing depth has three significant peaks at 1/2; **B.** *A. sinensis* has four significant peaks of minor allele frequency at 1/8, 1/4, 1/2, and 7/8, respectively, and SNP site sequencing depth has three significant peaks at 1/4, 1/2, and 7/8.

### Ploidy composition analysis

To further assess the ploidy composition of *A. sinensis*, we performed Smudgeplots analyses based on the depth and frequency of heterozygous *k*-mer pairs (with only one nucleotide difference, presented as A and B symbols) using the BGISEQ data from the normal reproductive animal. These analyses revealed that *A. sinensis* possesses an extremely complex ploidy composition, containing 41% of four octovalent (8n) *k*-mers (AAAAAAAB, AAAAAABB, AAAAABBB, and AAAABBBB), 3% of a hexavalent *k*-mer (AAAABB), 52% of two tetravalent (4n) *k*-mers (AAAB and AABB), and 4% a divalent *k*-mer (AB) (**Figure 3**A). The high proportion of octovalent *k*-mers demonstrated that the *A. sinensis* genome had typical octoploid characteristics. Moreover, we compared this analysis of *A. sinensis* with the results of four representative and well-studied tetraploid species – *Thymallus arcticus* (rediploidized autotetraploid) [35], *A. ruthenus* (rediploidized autotetraploid) [11], *Medicago sativa* (recently duplicated autotetraploid) [36], and *C. carpio* (allotetraploid) [37, 38] – using the Smudgeplots. *T. arcticus* and *A. ruthenus*, both rediploidized autotetraploids, show common characteristics whereby AABB accounts for the dominant proportion, and a lower and nearly equivalent proportion of AB. The proportion of AAAB in *T. arcticus* (3%) (**Figure 3**B) is much lower than in *A. ruthenus* (20%) (**Figure 3**C), which suggests that *T. arcticus* has a higher extent of rediploidization than *A. ruthenus* due to the lower evolution rate of Acipenseriformes, consistent previous research [11]. In *M. sativa*, AAAB accounts for the dominant proportion (63%), whereas AABB only accounts for 15% (**Figure 3**D), suggesting the more recent WGD of *M. sativa* (∼58 MYA) [36] in comparison to *T. arcticus* (80∼100 MYA)[35] and *A. ruthenus* (∼180 MYA) [11] resulted in a higher homology and lower rediploidization level. It is well known that *C. carpio* is one of the representative allotetraploid teleosts. Two ancient progenitor species (AA and BB) of *C. carpio* diverged 23 MYA, and they independently survived to subsequently produce the current allotetraploid *C. carpio* (AABB) by hybridization ∼12.3 MYA [37, 38]. Ploidy analysis of *C. carpio* shows that AB, AABB, and AAAB *k*-mers account for 59%, 30%, and 3%, respectively (**Figure 3**E). The distribution of heterozygous *k*-mers in the four tetraploid species showed that AB was the dominant proportion in allotetraploid species (*C. carpio*). This is due to the recombination of significantly differential subgenomes leading to *k*-mer pair sequences mispairing to tetravalents. Our results suggested that AAAB accounted for the dominant composition in the autotetraploid with lower rediploidization (*M. sativa*), whereas AABB was the dominant composition in autotetraploid with high rediploidization (*T. arcticus* and *A. ruthenus*) (**Figure 3**F). Compared with the tetraploid *A. ruthenus*, the octoploid *A. sinensis* may have undergone an additional round of WGD, resulting in the overlap of the genotypes we observe. Furthermore, the proportion of autopolyploidization characteristics, including AAAAAAAB, AAAAAABB, AAAAABBB, and AAAB, is up to 62% in *A. sinensis*, suggesting that the species is an autooctoploid.

**Figure 3.**
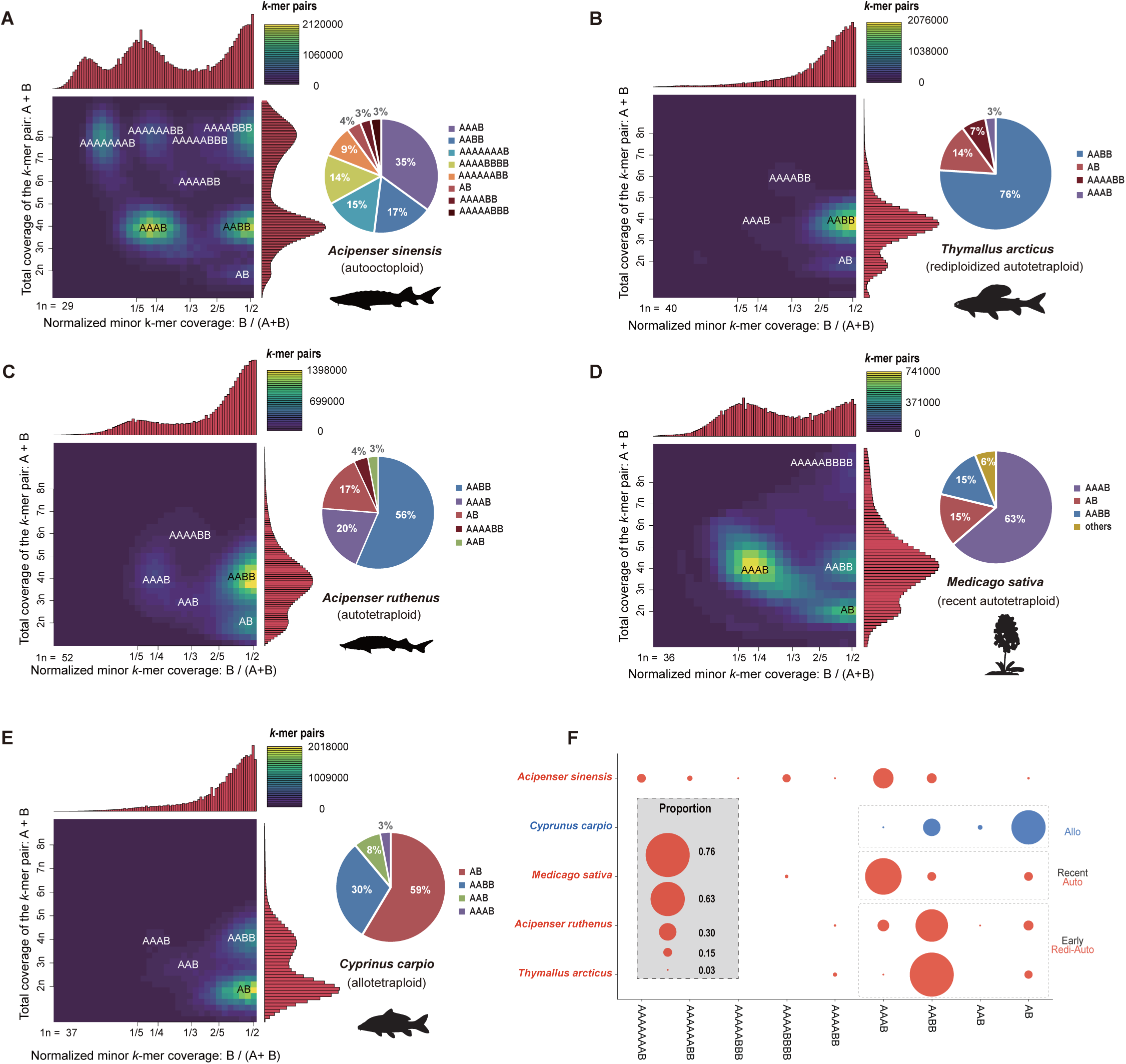
Ploidy compositions of five polyploidy species. **A.** Ploidy compositions of *Acipenser sinensis*. **B.** Ploidy compositions of *Thymallus arcticus***. C.** Ploidy compositions of *Acipenser ruthenus*. **D.** Ploidy compositions of *Medicago sativa***. E.** Ploidy compositions of *Cyprinus carpio*. The letters A and B in the Smudgeplots of the five polyploid species represent a pair of heterozygous *k*-mers with only one SNP difference. The brightness of each smudge is determined by the number of heterozygous *k*-mer pairs that fall within it. The percentage of each genotype is presented in the pie chart on the right. **F.** The distribution of nine genotypes in the five species. The main compositions of tetraploid species are framed by dashed lines. Allo, Recent Auto, and Early Redi-Auto represent allotetraploidization, recent autoploidization with low rediploidization, and early autoploidization with high rediploidization, respectively. AB (right composition) accounts for a higher proportion in allotetraploid, while AAAB and AABB (left composition) are higher proportions in autotetraploid. With increasing rediploidization, the proportion of left AAAB decreased while the right AABB increased.

Some repeats may be expanded specifically at each progenitor of subgenomes due to independent evolution before allotetraploid hybrids. Thus, the burst of distinctive TEs might be closely related to separating from two subgenomes in the allopolyploidy genomes [39-42]. The burst of distinctive transposable elements (TEs) might be closely related to the separate evolution of ancestors from two subgenomes in the allopolyploidy genomes [39-42]. Here, we struggled to detect the distinctive TEs between homologous sequences to identify the auto- or allo-polyploidization process of *A. sinensis*. However, no significantly specific TEs were observed (Figure S9; Tables S16 and S17), excluding the possibility of allooctoploidy, which was similar to what has been concluded for *A. ruthenus* [11]. These results implied that *A. sinensis* had undergone homologous duplications, i.e., is an autoocotoploid species.

### Timing of WGD and species divergence

To uncover a more accurate WGD and divergence timing for *A. sinensis* based on LORe and AORe, we screened 1,438 gene families with collinearity among the genomes of *A. sinensis* (S), *A. ruthenus* (R), *P. spathula* (P), and *Lepisosteus osseus* with the gene copy number of 2:2:2:1. We constructed 1,438 topologies using the screened gene families, and three types of representative topologies that accurately represented the AORe (PSR-PSR for topology name) and LORe (PP-SR-SR and PP-SS-RR for topology names) model (**Figure 4**A; Figure S1B) were collected (736 gene families). The PSR-PSR type was the dominant topology and accounted for 61.3% of 736 screened gene families, which was followed by PP-SR-SR (31.9%) and PP-SS-RR (6.8%) (Table S18). The PSR-PSR type was the most abundant in all of the topologies, indicating that the three species underwent a common duplication event, as otherwise this observation cannot be explained (Figure S1B). In addition, the PP-SR-SR ratio is the highest (31.9%), implying that the two families of Acipenseriformes diverged after the Acipenseriforme-specific WGD (As3R) but before complete rediploidization. Thus, LORe occurred during the evolution of Acipenseriformes species (Figure S1B). The low percentage of PP-SS-RR suggests that there was a nearly complete rediploidization event in the ancestors of *A. sinensis* and *A. ruthenus* before their divergence and speciation (Figure S1B). Allotetraploids are expected to show disomic inheritance (genetic diploidy) as soon as they are formed and the rediploidization is immediately completed [42]. As a result, high levels of LORe only appear in autopolyploid. Our results imply that the Acipenseriformes species shared a common polyploidization process. In addition, the distribution of LORe and AORe on the *A. sinensis*, *A. ruthenus*, and *P. spathula* chromosomes showed that AORe was mainly distributed on the 1-6 macro-chromosomes, whereas LORe tended to occur on medium- and micro-chromosomes, probably due to their instability (**Figure 4**B-D; Table S19). The synonymous substitution values (Ks) on the macro-chromosomes were larger than those on the medium- and micro-chromosomes (Figure S10). This supported the distribution features of AORe and LORe on the chromosomes.

**Figure 4.**
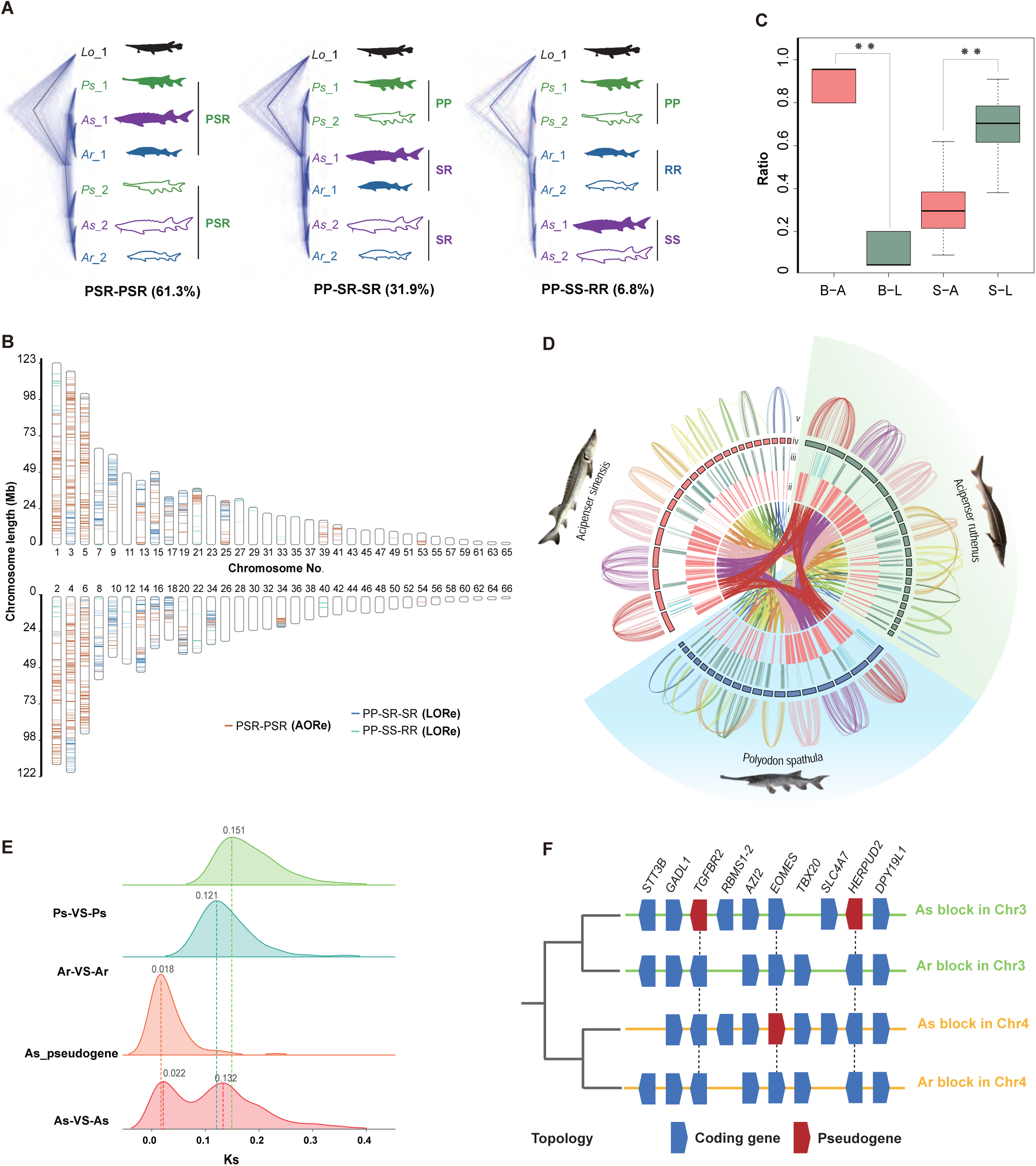
WGD events and divergence of Acipenseriformes based on analyses of LORe and AORe. **A.** Main topological structures based on different assumptions. L, P, S, and R represent *Lepisosteus osseus*, *Polyodon spathula*, *Acipenser sinensis*, and *Acipenser ruthenus*, respectively. SRP-SRP indicates that WGD, whole genome duplicaiton occurred before the divergence of the three species, which occurred LORe divergence. The proportion of collinear ohnologue pairs is 61.3% (451/736). PP-SR-SR indicates that WGD occurred before the differentiation between *A. sinensis* and *A. ruthenus* while following the divergence of *P. spathula* from Acipenseriformes and *P. spathula* has LORe differentiation. The proportion of collinear ohnologue pairs is 31.9% (235/736). PP-SS-RR indicates that the three species occurred LORe differentiation. The proportion of collinear pairs is 6.8% (50/736). **B.** Distributions of AORe and LORe on chromosomes. The red, blue, and green lines represent AORe in PSR-PSR, LORe in PP-SR-SR, and LORe in PP-SS-RR, respectively. **C.** Distributions of AORe and LORe homology on Macro- and Micro-chromosomes. B-A, B-L, S-A, and S-L represent the ratio of AORe on Macro-chromosomes, LORe on Macro-chromosomes, AORe on Micro-chromosomes and LORe on Micro-chromosomes, respectively. **D.** Distributions of AORe and LORe in the genome of *A. sinensis*, *A. ruthenus* and *P. spathula*. Tracks from the inner to the outside correspond to i) collinearity among *A. sinensis*, *A. ruthenus*, and *P. spathula*, ii) locations of AORe genes, iii) locations of LORe genes, iv) chromosomes with more than 20 genes (red, green and blue blocks represent chromosomes of *A. sinensis*, *A. ruthenus*, and *P. spathula*, respectively), and v) collinearity within each species. **E.** Estimation of WGD and divergence events occurred among *A. sinensis*, *A. ruthenus*, and *P. spathula* by synonymous substitution values (Ks). Ps, As, and Ar represent *P. spathula*, *A. sinensis*, and *A. ruthenus*, respectively. As3R is the first Acipenseriforme-specific WGD and Ass4R is the second *A. sinensis*-specific WGD. **F.** An example of coding genes and pseudogenes within one block in subgenomes-like of *A. sinensis* and *A. ruthenus*. It demonstrates that there are three pseudogenes (red) in *A. sinensis*.

Analyses of synonymous (Ks) substitution rates in coding genes and unitary pseudogenes were applied in estimating the timing of the *A. sinensis*-specific WGD (Ass4R) (Figure S11). Using As3R (T, 210.7 MYA; Ks rate, 0.132) as a reference, Ass4R (Ks rate, 0.022) was calculated at ∼35.12 MYA (**Figure 4**E). Furthermore, referring to previous studies, we speculated that some missing homoeologues, which were not detected as coding genes, would be presented in the *A. sinensis* genome as new unitary pseudogenes following Ass4R. We screened out 344 pseudogenes with high assurance from the gene families (with quadrivalent pairing collinearity in *A. ruthenus* and *A. sinensis* and each gene family conforming to the AORe model) **(Figure 4**F; Figure S12). Based on the accumulation of more non-synonymous (Ka) mutations than expected [42, 43], we estimated that most of these pseudogenes escaped evolutionary constraint ∼28.7 MYA (**Figure 4**E), which is in line with the expectation that they occurred shortly after Ass4R. Based on calculated Ks rates of three intra-species in Acipenseriformes (S: 0.132, R:0.121, and P:0.151), with common WGD time (T, 210.7 MYA), the absolute substitution rates of the sturgeons were 3.13×10^-10^, 2.87×10^-10^, and 3.58×10^-10^ per year, respectively, as calculated by the formula: Ks/(2T). *P. spathula* has the highest absolute substitution rate, followed by *A. sinensis* and *A. ruthenus*.

LORe cannot accurately reflect WGD due to delayed differentiation of LORe, whereas, AORe is able to better reflect the evolutionary trajectory of Acipenseriformes. The phylogenetic tree using MCMCTree analysis based on protein sequences of AORe in Acipenseriformes and other species ohnologues showed that the Acipenseriforme-specific common WGD (As3R) occurred 210.7 MYA. Divergence of paddlefish and sturgeon occurred ∼150 MYA. *A. sinensis* and *A. ruthenus* diverged 89.5–85.3 MYA (**Figure 5**), which is slightly later than the 121.3 (76.7–166.2) MYA estimated by a previous report based on a mitochondrial genome sequence data set of Acipenseriformes [33].

**Figure 5.**
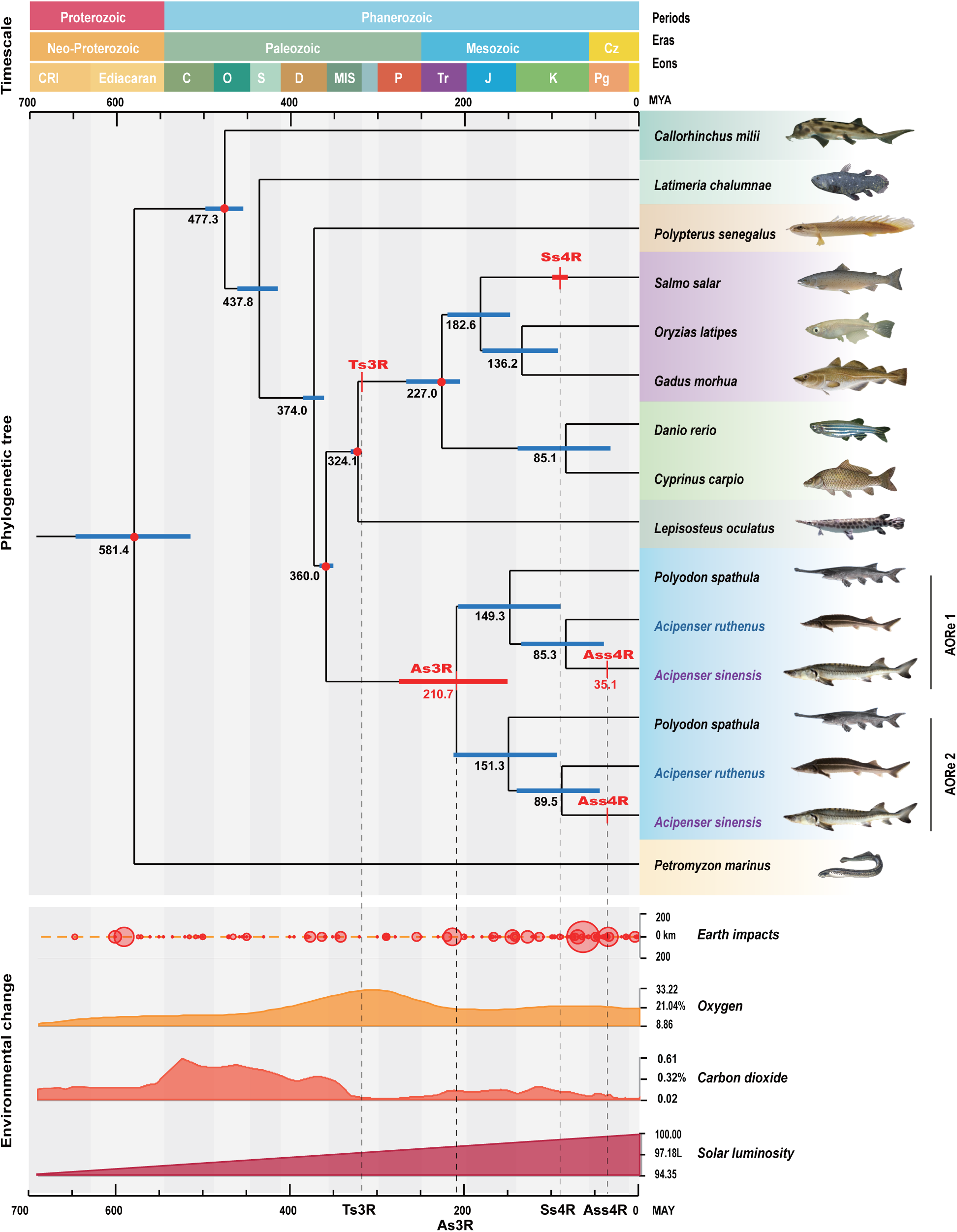
**Corresponding relationship among phylogenetic tree, whole genome duplication (WGD) events, and environmental changes.** Three red nodes are the intrinsic parameters based on fossil records. Black number indicates the divergence time. Blue block is the interval of estimated time. Red block indicates WGD time. Ts3R, Ss4R, As3R, and Ass4R are the teleost-specific 3^rd^ vertebrate WGD, the salmonids-specific 4^th^ vertebrate WGD, the Acipenseriforme-specific 3^rd^ vertebrate WGD, and the *Acipenser sinensis*-specific 4^th^ WGD, respectively. Environmental changes display asteroid impacts, oxygen, carbon dioxide, and solar luminosity in history.

## Discussion

The absence of whole-genome information has hindered the study of *A. sinensis* genetics and evolution, both of which could yield information aiding in conservation efforts and on fish ploidy evolution more broadly. *de novo* genome assembly of *A. sinensis* has been challenging because of the large size, higher polyploidy level, higher chromosome number, and complex chromosomal composition. In this study, we have successfully obtained a high-quality chromosome-level assembly of the *A. sinensis* genome. This is the first sequenced octoploid sturgeon genome in the order of Acipenseriformes and also the first sequenced genome of an octoploid vertebrate to date.

From our work presented here, the assembled genome size of *A. sinensis* is equal to the haploid genome size of *A. ruthenus* [11] and *P. spathula* [26]. The Hi-C anchoring rate of *A. sinenis* (98.3%) in this study is higher than that of *A. ruthenus* (90.2%) and *P. spathula* (96.5%) (**Table 1**). Moreover, high coverage of the complete genome indicates high integrity of the genome assembly. Genome completeness assessment shows that the proportion of complete BUSCOs (95.6%) is higher than *P. spathula* (93.7%) [26], *S. salar* genome (90.12% [44], *C. carpio* genome (81.70%) [37, 38], and the first *A. ruthenus* genome (81.6%) [30], slightly lower than the second *A. ruthenus* genome (98.3%) [11]. In addition, all chromosomes, especially the six macro-chromosomes, have high collinearity with *A. ruthenus* and *P. spathula*. Compared with recently reported high-quality genome assemblies of polyploid fishes, the quality of our *A. sinensis* genome assembly (contig N50: 4.06 Mb) has higher assembly quality as measured by contig N50 in comparison to the published tetraploid genomes of *A. ruthenus* (contig N50: 597.52 Kb) [11] and *P. spathula* (contig N50: 3.44 Mb) [26], and is close to that of hexaploid Prussian carp *Carassius gibelio* (contig N50: 4.3 Mb) [40]. According to the common diploid genome assembly, 132 chromosomes are theoretically supposed to be assembled for *A. sinensi*s. However, the homologous copies of each chromosome result in the collapse of polyploid genome assemblies and thus form a “mosaic” reference from both parental haplotypes as a “monoploid” representation of the genome. For the autooctoploid *A. sinensis*, we were able to construct 66 chromosomes based on the final assembly. This includes 6 macro-chromosomes, 60 medium-chromosomes, and micro-chromosomes, which corresponded to 1/4 of the total number of chromosomes or two monoploids of this species. Overall, while we will of course focus future work on reconstructing the remaining chromosomes, which presented significant challenges due to the complexity of the genome, we consider our current genome assembly of *A. sinensis* to be of high quality for an octoploid species with such a high DNA content and complex chromosome compositions.

Whole genome sequencing provides a powerful solution for revealing ploidy composition and evolution. Smudgeplot analyses on basis of our whole genome sequencing data showed that *A. sinenes* and *A. ruthenus* ploidy are not well-defined octoploid or tetraploid as previously thought. Rather, they exhibit a complex transition ploidy with multiple compositions arising from octoploid rediploidization. We observed that *A. ruthenus* has five different ploidy compositions and propose that the species is tetraploid with a certain degree of rediploidization, which is consistent with previous studies of segmental rediploidization [11]. In contrast, we find that *A. sinensis* has eight more complex ploidy compositions than *A. ruthenus* and should therefore be considered a paleooctoploid experiencing diploidization. These results are consistent with our SNP analyses in this study and previous reports using cytogenetic and molecular methods [18, 22-24]. These conclusions potentially supports the latest viewpoint that ploidy groups A and B are evolutionary tetraploid and octoploid, respectively [17-19].

We further explored whether *A. sinensis* and *A. ruthenus* underwent homologous or heterologous polyploidization, respectively. Based on the Smudgeplot analyses of four representative tetraploid species, we propose a criterion for distinguishing between homologous and heterologous WGDs. Our results showed that *A. sinensis* and *A. ruthenus* have striking autopolyploid characteristics based on this criterion. Burst of distinctive TEs might be closely related to separate from two subgenomes in the allopolyploidy genomes. We thus also carried out a comparison of the TE landscape of *A. sinensis* paralogous chromosomes to explore whether *A. sinensis* was homologous or heterologous. For allopolyploidy, the fast-evolving repeats and relics of mobile elements are specific to their allopolyploid ancestors and thus have significant differences, whereas for an autopolyploid chromosome set, the repeat elements would not differentiate the homologs and thus the individual TE families in paralogous chromosomes were monophyletic [11, 26]. We did not detect significantly specific TE families in the *A. sinensis* genome in this study. Our results were similar to the analysis of the *A. ruthenus* genome [11] and therefore, we conclude that *A. sinensis* is likely to be an autooctoploid.

Phylogenetic trees have shown that the Acipesneriforme ancestor diverged from the teleost ancestor ∼360 MYA, which is earlier than the teleost-specific WGD (Ts3R) occurring approximately 320–350 MYA and thereby implying that Acipenseriformes experienced an independent WGD [45]. In previous reports, WGD and divergence times of Acipenseriformes were calculated based on single-copy genes or homolog genes by whole genome sequencing. For example, Cheng et al. reported that paddlefish (Polyodontidae) and sturgeon (Acipenseridae) diverged around 81.5 MYA and a round of WGD event in the American paddlefish occurred about 46.6–54.1 MYA, implying that this WGD event independently occurred in the American paddlefish after the species diverged [26]. In contrast, Du et al. reported that the *A. ruthenus* WGD occurred around 180 MYA [11]. We feel that it is more reasonable to calculate WGD and divergence time only based on AORe to eliminate the LORe interference. Integrating AORe, our results show that Acipenseriformes shared a common WGD event (As3R) dating back to 210.7 MYA before the divergence ∼150 MYA (149.3–151.3 MYA) of Acipenseridae and Polyodontidae, and *A. sinensis* underwent an additional lineage-specific WGD (Ass4R) around 35.12 MYA, which resulted in the speciation of an autooctoploid. Interestingly, a similar study on WGD based on LORe and AORe using *A. ruthenus* and *P. spathula* genome was performed by Redmond *et al.*[46] In this work, they screened a total of 5,439 gene families containing high confidence ohnolog pairs in two species, *A. ruthenus* and *P. spathula* to analyze maximum likelihood gene trees. In comparison, we screened 1,438 gene families from 4 species including *A. sinensis*, *A. ruthenus*, *P. spathula*, and *L. osseus*. Their study excluded independent WGDs and found a high proportion of tetraploidy (∼50-66% of the genome) at the time of speciation, which is inconsistent with past studies inferring independent WGD events [26, 30, 47-49]. Consistent with our estimates of WGD timing, Redmond *et al*. inferred a divergence time of ∼171.6 MYA (95% credibility interval range: ∼124.1–203.3 MYA) for the split of sturgeons and paddlefish, and estimated a lower bound for the shared sturgeon-paddlefish WGD (As3R) at ∼254.7 MYA (95% credibility interval range: ∼207.1–289 MYA) [50]. Excitingly, the As3R time is close to the report (∼180 MYA) by Du *et al*. [11], and the divergence time is consistent with the estimated time (141.4 MYA) based on mitochondrial genome sequences data sets by Peng et al. [33]. Thus, we believe that the WGD and divergence time reported here are particularly robust. Analyses based on AORe of excluded LORe interference therefore provides a novel method for calculating more reasonable WGD and divergence time for polyploid species.

WGD events were strongly correlated with the timing of drastic environmental changes [51], such as Earth impacts and dramatic changes in oxygen concentration, carbon dioxide concentration, or temperature [52]. Interestingly, the larger events of Earth impacts in geological history correspond well to As3R and Ass4R in this study (**Figure 5**), suggesting that the violently climatic and geological changes caused by Earth impacts potentially resulted in two rounds of WGDs by affecting the reproductive process of *A. sinensis* ancestors.

Overall in this study, we have accomplished the whole genome sequencing of the first octoploid fish and revealed the specific ploidy composition and WGD evolutionary history of *A. sinensis*. This high-quality genome resource will serve as a powerful platform for the studies of genetics, evolution, and conservation in Acipenseriformes species, as well as provide a reference for the genomic studies of other polyploid vertebrates.

## Materials and methods

Complete description of Materials and Methods can be found in the File S1 Supplementary information.

### *A. sinensis* animals used in study

All *A. sinensis* used in this study were derived from an artificially bred stock. A male gynogenic *A. sinensis* was used for genome sequencing and assembly. An animal that was derived through sexual reproduction and normal development rather than meiotic gynogenesis, was sampled for ploidy estimation and transcriptome sequencing. The experiments were performed in according with the guidelines of the Institutional Review Board on Bioethics and Biosafety of the Chinese Sturgeon Research Institute.

### Genome and transcriptome sequencing

The DNA for sequencing was extracted from the blood of a diploid meiotic gynogenetic progeny of *A. sinensis* (male, 3 years old) (Figure S13). The short reads were sequenced using three paired-end (PE) libraries (170 bp, 500 bp, and 800 bp) using the Illumina HiSeq 2000 platform. We applied rigorous criteria to filter the raw reads generated by PE libraries into clean reads using SOAPfilter in the SOAPdenovo package [53, 54]. The single-molecular long reads were sequenced using eight libraries using the PacBio Sequel sequencing platform. A total of 42 SMRT Cells were sequenced using the 20 Kb large insert size libraries for genome assembly. The genomic large DNA from the fresh blood sample of *A. sinensis* was used for the Hi-C library construction. The extracted DNA with a size of 300-350 bp was sequenced on a BGISEQ-500 sequencing platform.

A mixed sample containing 11 different tissues was sequenced using the PacBio Sequel sequencing platform. In addition, nine samples of three different tissues, containing the hypothalamus, pituitary, and gonad, from 3 female control individuals (normally sexual reproduction) were used for RNA sequencing by the Illumina platform [55].

### Genome size estimation

The genome size of *A. sinensis* was estimated by flow cytometry (FCM) of red blood cells from a normal reproductive animal. Three different *k*-mers (19-, 21-, and 23-mer) analyses were also implemented using Jellyfish [56] by genomic clean reads within small insert size libraries to predict the genome size. The total genome size was estimated according to the following formula: genome size = *k*-mer number/peak depth, where *k*-mer number is the total number of *k*-mers, and peak depth is the maximal frequency.

### Genome assembly and chromosome anchoring

PacBio sequencing raw data were corrected using Canu (https://github.com/marbl/canu) software. The corrected PacBio reads were assembled into original contigs using Smartdenovo (https://github.com/ruanjue/smartdenovo). The original contigs were corrected and polished using Arrow [57] and Pilon [58] with PacBio data and Illumina HiSeq data, respectively. Furthermore, PurgeDups (https://github.com/dfguan/purge_dups) was implemented to break misjoins and generate a final assembly. The contigs were then anchored to chromosomes using Juicer (https://github.com/aidenlab/juicer) and 3D-DNA (https://github.com/aidenlab/3d-dna) pipeline with the Hi-C data. Smartdenovo has demonstrated notable effectiveness in handling polyploid and highly heterozygous genome assemblies. Arrow and Pilon are capable of effectively correcting erroneous nucleotide bases. PurgeDups is also a commonly employed method for filtering redundant sequences and obtaining haploid assemblies.

### Genome annotation

We predicted genes in the genome of *A. sinensis* using ab initio-based, homology-based and transcriptome-assisted methods. De novo gene prediction was performed using AUGUST and SNAP. The protein sequences of *Callorhinchus_milii*, *Danio_rerio*, *Latimeria_chalumnae*, *Lepisosteus_oculatus*, and *Petromyzon_marinus* (Ensembl-100 version) were downloaded from the Ensembl database for homology-based gene set prediction by Exonerate software. Gene structures were annotated using three approaches (ab initio predictions, homolog proteins, and transcriptome data) that were combined using MAKER software. Furthermore, gene functions were annotated against seven public databases including the NCBI non-redundant protein sequences (Nr), Swiss-Prot, Kyoto Encyclopedia of Genes and Genomes (KEGG), Cluster of Orthologous Groups of proteins (COG), TrEMBL, InterPro and GO databases according to the best match of the alignments using blastp (E-value < 1×10^-5^).

Two kinds of repeats, tandem repeats and transposable elements (TEs), were identified before performing genome annotation. Tandem repeats were predicted using Tandem Repeats Finder (v4.09) [59]. TEs were detected based on homolog and de novo strategies. For the homolog approach, TEs were predicted using RepeatMasker [60] and RepeatProteinMask based on the Repbase database [61] and the TEs database in the RepeatMasker software packages, respectively. For the de novo approach, the de novo repeat library was predicted using RepeatModeler (RepeatModeler-open-1.0.11) (http://www.repeatmasker.org/RepeatModeler/), and TEs were annotated by RepeatMasker software based on the de novo library.

### Collinearity analyses, gene family identification, and phylogenetic tree construction

MCscan (Python version) [62] was used for the genomic analysis between *A. sinensis*, *Atractosteus spatula*, *Acipenser ruthenus*, and *Psephurus gladius*. The collinearity figure was drawn based on the collinear gene pairs information between species.

Thirteen species (*Acipenser sinesis*, *Acipenser ruthenus*, *Callorhinchus milii*, *Danio rerio*, *Gadus morhua*, *Latimeria chalumnae*, *Lepisosteus oculatus*, *Oryzias latipes*, *Petromyzon marinus*, *Polyodon spathula*, *Polypterus senegalus*, *Cyprinus carpio*, and *Salmo salar*) were used in the phylogenetic analysis. The protein-coding genes of these species were downloaded and filtered, and only the longest open reading frame (ORF) with a gene encoding more than 50 amino acids remained for the gene family clustering and phylogenetic analysis. Because rediploidization of *P. spathula*, *A. sinesis*, and *A. ruthenus*, the protein-coding genes were separated into two haplotype sets in the following analysis process. Gene families were identified with Orthofinder [63]. The single-copy orthologous genes from gene families were further aligned using MUSCLE (MUSCLE, RRID: SCR_011812, version 3.8.31) [64] with default parameters and subsequently reverse-translated into codon sequences. These aligned sequences were concatenated to generate a super alignment matrix for phylogenetic reconstruction based on (PhyML, RRID: SCR_014629) [31] with four-fold degenerate (4D) sites of the single-copy orthologs shared among the 13 species. Additionally, we employed IQ-TREE (v1.6.12) [65] to construct gene phylogenetic trees of the single-copy orthologs and used Astral (v5.6.1) (https://github.com/maryamrabiee/Constrained-search) to integrate the gene trees. The resulting phylogenetic tree was consistent with the tree generated by PhyML. The divergence time was determined by MCMCTree from the PAML version 4.5 package (PAML, RRID: SCR_014932) [66, 67] with the approximate likelihood calculation method, the ‘correlated molecular clock’ and ‘REV’ substitution model, successively. Three divergence dates from the TimeTree database [68] were used for calibration.

### Ploidy evaluation

Ploidy of the species was estimated by the maximum number of alleles per individual at each microsatellite loci. To obtain available simple sequence repeat (SSR) markers for determining the ploidy of *A. sinensis*, all screened SSRs of a tetra-nucleotide repeat were verified by polyacrylamide gel electrophoresis (PAGE) and capillary electrophoresis on the ABI 3730 Genetic analyzer, respectively.

Paired-end reads of *A. sinensis* and *A. ruthenus* were mapped to their assembled scaffolds by aligner BWA (Version 0.7.12-r1039) and Samtools (Version 1.4). The heterozygous SNPs were called by Freebayes (v0.9.10-3-g47a713e). The average allele mapping depth and the minor allele frequency of the variant sites were calculated to estimate ploidy based on the heterozygous sequence polymorphism.

### Heterozygous *k*-mer pairs analysis

To disentangle the genomic ploidy of *C. carpio*, *T. arcticus*, *M. sativa*, *A. sinensis*, and *A. ruthenus*, we extracted the haplotype structures from heterozygous *k*-mer pairs by using the Smudgeplots pipeline [69]. First, we produced a *k*-mer frequency file by KMC [70] with k = 21 from trimmed reads. Then, we searched for all heterozygous *k*-mer pairs that differed at exactly one nucleotide through a systematic scan of all input *k*-mers. To avoid sequencing errors with genomic *k*-mers, we filtered the *k*-mers with a depth of less than 14, which was the depth of the first trough in the *k*-mer frequency curve. Finally, we performed the R script of the pipeline, plotted the Smudgeplot, and estimated ploidy using the coverage file of heterozygous *k*-mers. This tool performed gymnastics with the heterozygous *k*-mer pairs by comparing the sum of *k*-mer pair coverages (CovA + CovB) to their relative coverage (CovA / (CovA + CovB)).

### LORe and AORe analysis

Combined with gene family identification and genome collinearity analysis, we identified the potential ohnologues with 2:2:2:1 in *A. sinesis*, *A. ruthenus*, *P. spathula*, and *L. oculatus.* The protein sequences of ohnologues were aligned using MUSCLE (v3.8.425) [64] with the default parameters. These alignments were subsequently converted into coding sequence (CDS) alignment by tracing the coding relationship using pal2nal.v14 [71]. Gblocks (v0.91b) [72] was employed to conduct further checks (trim) on the CDS alignments with parameters “-t = c”. The trimmed alignments with lengths less than 150 bp were filtered and then transmitted to IQ-Tree (v1.6.12) [65] to infer the gene tree with settings: -alrt 1000 -bb 1000. Each trimmed gene was subjected to a gene tree analysis in the same manner. DensiTree [73] was used to visualize the topologies of these trees.

The protein sequences of AORe ohnologues were selected. The ohnologues of AORe exhibited a pairable topology of PSR-PSR, as inferred from the aforementioned three species analysis. The protein sequences of AORe ohnologues were aligned using MUSCLE (v3.8) [64] with default parameters, and subsequently reverse-translated into codon sequences. These aligned sequences were concatenated to generate a super alignment matrix, and phylogenetic reconstruction was performed using the maximum-likelihood (ML) method in IQ-TREE (v1.6.12) [65]. The best-fit evolutionary substitution model was determined using ModelFinder. Basing on the phylogenetic tree, the divergence time between individual species and ohnologue subgenomes was estimated using MCMCTree by using the proteins with the approximate likelihood calculation method, the ‘correlated molecular clock’, and ‘REV’ substitution model, successively. Four data from the TimeTree database [68] were used for calibration. We calculated Ks of AORe ohnologues of inter-species and intra-species in the three genomes. Ks analyses were performed using default parameters and the ‘fasttree’ node-weighting method in the wgd package [31, 64, 66, 74]. Log normal distributions in Ks were plotted based on node-averaged values as calculated in the wgd package. The Gaussian mixture models (GMMs) were fitted to the Ks distribution following the wgd pipeline, with the optimal number of components assessed using the Bayesian information criterion.

## Data availability

The assembled genome sequences and gene annotations have been deposited in the Genome Warehouse database at the National Genomics Data Center, Beijing Institute of Genomics, Chinese Academy of Sciences / China National Center for Bioinformation (GWH: GWHBQEF00000000; Reviewer link: https://ngdc.cncb.ac.cn/gwh/Assembly/reviewer/IHkWasXWrgtQyrZBJcLVAqIJzIhTTlLCCghTRvWRnZSNKtYQTjBawrLMgyPYCksc). The raw datasets for genome assembly have been deposited in the Genome Sequence Archive at the National Genomics Data Center, Beijing Institute of Genomics, Chinese Academy of Sciences / China National Center for Bioinformation (GSA: CRA009603, Reviewer link: https://ngdc.cncb.ac.cn/gsa/s/Ovxx4S42; CRA009438, Reviewer link: https://ngdc.cncb.ac.cn/gsa/s/4FXIUZ6Z).

## Author contributions

**Hejun Du**, **Lei Chen**, **Binzhong Wang**, and **Xueqing Liu:** presided over the project application, including experiments design, project management, sturgeon breeding, sample preparation, research design, data collection, and manuscript co-drafting and modification. **Binzhong Wang**, **Bin Wu**, and **Jianbo Jian:** performed study design, data input and processing, genome assembly and annotation, manuscript co-drafting, and modification. **Binzhong Wang**, **Bin Wu**, **Yao Ming,** and **Yanhong Li:** performed the data input and processing, genome assembly and annotation, evolution analysis, and manuscript drafting. **Mingzhou Bai**, **Juanjuan Liu**, **Qingkai Zeng,** and **Hongqi Wang:** data output, project management, and sample preparation. **Kan Xiao**, **Yacheng Hu**, **Baifu Guo**, **Chun Tan**, and **Xun Zhao:** bred and cultivated the sample for genome and transcriptome sequencing. **Jing Yang**, **Zhen Yue**, **Zixuan Hu**, **Junpu Mei**, and **Yong Gao:** revised the manuscript and discussed the data. **Lei Chen**, **Zhiyuan Li**, **Yuanjin Yang**, and **Wei Jiang:** presided over the project application, supervised the study, and revised the manuscript. All authors provided comments and suggestions for improvement of the manuscript.

## Acknowledgments

This work was supported by the Three Gorges Environmental Funds of China Three Gorges Corporation (XN270). We acknowledge the assistance from Debin Shu, Jianming Zhang, Jianyi Wan, Jiayuan Tian, Hua Jiang, Jun Rao, and other colleagues for the sample preparation. We used the services of Life Science Editors for editing of the manuscript. We very gratefully thank Professor Jianfang Gui and Professor Shunping He, Institute of Hydrobiology, Chinese Academy of Sciences, Professor Liwu Lu, Beijing geological museum, and Professor Daqing Li, Gansu Agricultural University, for their valuable suggestions.

## Supplementary material

**File S1 Supplementary information**

**Figure S1 Relationships among WGD, divergence, and rediploidization based on AORe and LORe model.**

**A.** The AORe model of post-WGD, and the LORe model of post-WGD evolution following delayed rediploidization. **B.** PSR-PSR, PP-SR-SR, and PP-SS-RR model. P, S, and R represent Polyodon spathula, Acipenser sinensis, and Acipenser ruthenus, respectively. PSR-PSR was constructed by AORe and the divergence and speciation occurred after Acipenseriforme-specific WGD (As3R) and the complete rediploidization. PP-SR-SR was constructed by LORe and the two families of Acipenseriformes diverged after As3R before the complete rediploidization. PP-SS-RR was constructed by LORe and the divergence and speciation occurred before the complete rediploidization after As3R.

**Figure S2 19-mer, 21-mer, and 23-mer distributions were used for the estimation of genome size.**

The X-axis represents the depth of sequencing. The Y-axis is the proportion of 17-mer frequencies at different depths. The total genome size was estimated according to the following formula: genome size = *k*-mer number/peak depth, where *k*-mer number is the total number of *k*-mers, and peak depth is the maximal frequency.

**Figure S3 Flow Cytometer result of *Acipenser sinensis*.**

**Figure S4 Veen diagram of gene function annotation based on Nr, InterPro, KEGG, SwissProt, and KOG databases.**

A total of 22,939 genes were shared by the eight database annotations.

**Figure S5 Phylogenetic tree constructed with 2,096 single copy orthologous genes using Astral.**

**Figure S6 Phylogenetic tree constructed with 2,096 single copy orthologous genes using PhyML.**

The branch length represents the neutral divergence rate.

**Figure S7 Karyotype of *Acipenser sinensis*.**

A total of 264 chromosomes were identified (left) and ranged in groups containing four chromosomes (right). The first row is the macro-chromosomes.

**Figure S8 Ploidy identification based on SSRs.**

Eight peaks were identified by tetra-nucleotide repeat SSRs using capillary electrophoresis on ABI PRISM 3730 Genetic Analyzer.

**Figure S9 Detection of homology and heterology based on differential TEs screening.**

Pair_blocks_1-10 at the X-axis are 10 homoeologous sequence blocks with collinearity. The Y-axis represents 20 TEs in the 10 homoeologous blocks. Nsd represents that the block pair of TE has no significant difference. If at least one Nsd block appears in a row, the row is not a differential TE.

**Figure S10 Homologous gene dot plot within the *Acipenser sinensis* genome.**

Ks value for homologous genes in each inferred collinear block is shown. The high Ks values were presented on macrochromosomes 1-6.

**Figure S11 Distribution of Ks analysis in coding genes and unitary pseudogenes.**

**Figure S12 Amino acid sequence of the COG7 coding gene and pseudogene alignment in *Acipenser sinensis* and *Acipenser ruthenus*.**

Four genes were selected, As-Pseudogene1 (Gene id: ACSI009096-D2) and Ar-Gene1 (ACSI009096), As-Gene2 (XP_033886174.2) and Ar-Gene2 (XP_033907760.1) were from the common AORe, respectively. Missing codons are marked with dashes. Frameshifts and premature stops are marked by * and X, respectively (pointed by the red arrow and blue arrow).

**Figure S13 Identification results for gynogenetic *Acipenser sinensis* by microsatellite DNA analysis on PAGE.**

Lane M represents the DNA ladder marker. Lanes 1-21 show gynogenetic individuals. Lanes D and S represent the dam (maternal) and sire (paternal), respectively. Lanes C1-C5 represent the control diploid individuals.

**Table S1 Statistics of Acipenser sinensis sequence data derived from paired-end and mate-paired sequencing by Illumina platform.**

**Table S2 Raw data statistics of PacBio sequencing.**

**Table S3 Statistics of Hi-C sequencing and genome alignment.**

**Table S4 Statistics of chromosome relative length of *Acipener sinensis*.**

**Table S5 Statistics of genome size estimation by different k-mer analysis.**

**Table S6 Statistics of BUSCOs estimation of *Acipenser sinensis* genome assembly.**

**Table S7 Statistics of RNA-seq data.**

**Table S8 Statistics of gene annotation.**

**Table S9 Statistics of gene fuction annotation.**

**Table S10 General statistics of repeats in genome.**

**Table S11 Stactistics of TEs content in genome.**

**Table S12 Sythenic gene statistics in different species groups.**

**Table S13 Statistics of gene family analysis based on OrhoFinder.**

**Table S14 Distribution of tetra-nucleotide repeats (simple sequence repeats, SSRs) in screening process.**

**Table S15 Comparision of SNPs and heterozygosity.**

**Table S16 Homoeologous block with collinearity for specific repeats cluster.**

**Table S17 Statistics of TE family in the 10 block-pairs (bp/1Mb).**

**Table S18 Statistics of three topological structure based on LORe and AORe.**

**Table S19 Statistics of three topological structures in large chromosomes and small chromosomes of three Acipenseriforme species.**

