## Supplementary material for "Whole Genome Sequencing Reveals Autooctoploidy in the Chinese Sturgeon and its Evolutionary Trajectories": File S1 Supplementary information

#### Sampling preparation

*A. sinensis* in this study was artificially bred. Heterozygosity is a major challenge for efficient and high-quality genome assembly, especially for the complex genome of Chinese sturgeon. Here, to reduce the impact of heterozygosity and improve the quality of whole-genome assembly, meiotic gynogenetic progenies of *A. sinensis* (GP) were produced by heat-shock treatment after egg activation with ultraviolet-irradiated homologous sperm in 2013. Then, the progenies were confirmed as exclusively maternal inheritance by using microsatellite DNA analysis in 2014 (Figure S12). The blood from a normal individual (NI), which was the female parent of the meiotic gynogenetic *A. sinensis*, was collected for genome sequencing to reveal ploidy and ploidy composition. All of the blood was stored at -80 °C for DNA extraction.

Multiple tissue samples from normal individuals in the population were collected for transcriptome sequencing to assist genome annotation. A total of 11 *A. sinensis* were selected to sample. Nine samples of three different tissues, containing hypothalamus (H), pituitary (P) and gonad (G), and 11 tissues containing brain, gill, muscle, liver, spleen, kidney, heart, skin, stomach, testis, and ovary from NI were rapidly collected and flash frozen in liquid nitrogen, then stored at -80 °C for RNA extraction

In addition, 62 normal *A. sinensis* samples were used to screen microsatellite markers.

#### Genomic DNA and RNA preparation

Total genomic DNA was extracted from the blood of a male gynogenetic *A. sinensis* and its female parent using the Blood and Tissue Kit (Qiagen, Valencia, CA) following the manufacturer's instructions. Total RNA was extracted with Trizol reagent (Invitrogen) following the manufacturer's instructions. The RNA quality, integrity, and concentration were calculated and checked by Agilent 2100 Bioanalyzer (Agilent, Santa Clara, CA), and the high-quality RNA (RNA Integrity Number > 7.0) was used for transcriptome sequencing.

#### Library construction, sequencing and data processing

A total of three paired-end (PE) libraries (170 bp, 500 bp, and 800 bp) were constructed with acoustic shearing using a Covaris instrument (Covaris, Woburn, MA), and sequenced by Illumina HiSeq 2000 platform (Illumina, CA, USA) with sequence reads lengths: 100 bp. Eight libraries of

350 bp short insert were constructed and sequenced in 150 bp paired-end mode on the BGISEQ-500 platform. We applied rigorous criteria to filter the raw reads generated by PE libraries into clean reads using SOAPfilter in the SOAPdenovo package(Luo, et al. 2012; Luo, et al. 2015). First, we removed the raw reads containing an adapter ≥ 10 bp with a mismatch of ≤ 3 bp and > 1% N content or duplicated paired-end reads. Second, we filtered out the reads of more than 50% low-quality bases (Q-value ≤ 15). The PE reads were corrected using SOAPec (version 2.03)(Luo, et al. 2012; Luo, et al. 2015), a short-read correction tool that allows a selection of sets according to *k*-mer size.

Besides, the 20 Kb large insert size libraries were constructed and applied in genomic long-read sequencing using a blood sample of *A. sinensis*. First, the large fragment of genomic DNA was collected and concentrated by AMPure PacBio beads, then applied in SMRTbell preparation based on the manufacturer's instruction (Pacific Biosciences, CA, USA). Eight libraries were constructed and sequenced by the PacBio Sequel sequencing platform (Pacific Biosciences, CA, USA). In total, 42 SMRT Cells were sequenced and generated 221.96 Gb sequencing data for genome assembly.

The genomic large DNA from the fresh blood sample of *A. sinensis* was used for the Hi-C library construction. The extracted DNA was fragmented to a size of 300-350 bp and sequenced on a BGISEQ-500 sequencing platform. For Hi-C sequencing, blood cells were fixed with 2% formaldehyde for *A. sinensis*. The cross-linked DNA was digested with MboI and the sticky ends were biotinylated by incubating with biotin-14-dATP and Klenow enzyme. After DNA purification and removal of biotin from unligated ends, Hi-C products were enriched and physically sheared to fragment sizes of 200-300 bp. The biotin-tagged Hi-C DNA was pulled down and processed into paired-end sequencing libraries which were sequenced PE 100 bp on the BGISEQ-500 sequencing platform. Finally, 799 Gb Hi-C data were obtained.

Nine H, P, and G RNA samples were used for Illumina high-throughput RNA sequencing. First, total RNA samples (5 μg) were subjected to library construction for Illumina sequencing following the protocol of the mRNA sample preparation kit (Illumina). Then, the nine libraries were sequenced using Illumina HiSeqTM 2000 with 2 × 90 paired-end sequencing. To obtain high-quality reads, the raw reads were filtered by SOAPnuke (v1.5.6)(Chen, et al. 2017) with the default parameters except "-l 20 -q 0.2 -n 0.05". The full-length transcriptome was sequenced for genome annotation assistance by the PacBio Sequel sequencing platform (Pacific Biosciences, CA, USA) using mixed samples from 11 different tissues, and then processed by IsoSeq technology through the SMRT method (https://www.pacb.com/smrt-science/smrt-sequencing/) to obtain full-length non-chimeric reads.

**Genome assembly**

We first corrected the 221.96 Gb PacBio sequencing data using Canu (https://github.com/marbl/canu) software with default parameters. Afterward, the contig-level genome was assembled with Smartdenovo (-J 5000 for wtpre; -k 19 -z 10 -Z 16 -U -1 -m 0.1 -A 5000 for wtzmo; -d 3 -k 100 -m 0.1 -FT for wtclp; -w 300 -s 200 -m 0.6 -r 0.95 -c 1 for wtlay) (https://github.com/ruanjue/smartdenovo) using the corrected PacBio reads. The original contigs were then corrected and polished using Arrow(Chin, et al. 2013) and Pilon(Walker, et al. 2014) with PacBio data and Illumina HiSeq data from small insert size libraries, respectively. Furthermore, PurgeDups (https://github.com/dfguan/purge_dups) was implemented to break misjoins and generate a final assembly. Then scaffolding was performed using Hi-C based information, in which Hi-C reads were aligned to the assembled draft genome by Juicer v1.6 (https://github.com/aidenlab/juicer(Durand, et al. 2016)), and contigs were mapped onto chromosome-level scaffolds using 3D-DNA(Dudchenko, et al. 2017) using default parameters. Manual checking and refinement of the draft assembly were carried out via Juicebox Assembly Tools (https://github.com/aidenlab/Juicebox, v1.1108).

**Genome size estimation**

The genome size of *A. sinensis* was estimated by flow cytometry (FCM) of red blood cells from a normal individual. The reference is the blood cells of *Gallus gallus* (Detailed statistics see Figure S3). Meanwhile, *k*-mer analysis was implemented using Jellyfish(Marcais and Kingsford 2011) by genomic clean reads within small insert size libraries to predict the genome size. Here total *k*-mer numbers and peak depth were calculated by three different *k*-mers (19-, 21-, and 23-mer), respectively. The total genome size was estimated according to the following formula: genome size = *k*-mer number/peak depth, where *k*-mer number is the total number of *k*-mers, and peak depth is the maximal frequency.

Approximately 511.01 Gb raw sequencing data from short insert size libraries (170, 500, 800 bp libraries) was created by the Hiseq sequencing platform, and then 421.58 Gb filtered data were used to predict the genome size (Table S1). First, three different *k*-mers (19-mer, 21-mer, and 23-

mer) were used to estimate the genome size, and 21-mer was finally elected. Based on 21-mer frequencies, we calculated that peak depth was 173×, and the number of the 3 *k*-mers presented in this subset is 341,727,680,524 (Table S4). Then, the two monoploid genome sizes, which were estimated by dividing the total number of 21-mer using the third peak depth, were 1.975 Gb (Table S4). The 2C nuclear DNA content was also estimated by Flow Cytometer compared with Chicken (~2.2 pg/2C) (Figure S3). The results showed that the nuclear DNA content of *A. sinensis* is approximately 9.07 pg corresponding to four sets of sub-genomes. The two monoploid genome sizes will be approximately 2.21 Gb. In previous studies, sturgeons with ~240 chromosomes normally had a 2C nuclear DNA content of 8 ~ 9 pg(Blacklidge and Bidwell 1993; Zhang, et al. 1998; Bytyutskyy, et al. 2012). In this study, the estimated two monoploid genomes of 1.975 Gb, which is slightly less than the value of flow cytometry of approximately 2.21 Gb (Figure S2), were finally identified as the genome size of *A. sinensis*.

**Genome annotation**

Two kinds of repeats, tandem repeats and transposable elements (TEs), were identified before performing genome annotation. Tandem repeats were predicted using *Tandem Repeats Finder* (v4.09)(Benson 1999). Transposable elements (TEs) were detected based on homolog and *de novo* strategies. For the homolog approach, TEs were predicted by *RepeatMasker*(Tarailo-Graovac and Chen 2009) and *RepeatProteinMask* based on the *Repbase* database(Bao, et al. 2015) and the TEs database in the *RepeatMasker* software packages, respectively. For the *denovo* approach, the *denovo* repeat library was predicted by *RepeatModeler* (RepeatModeler-open-1.0.11) (http://www.repeatmasker.org/RepeatModeler/), and TEs were annotated by *RepeatMasker* software based on the *denovo* library.

We predicted genes in the genome of *A. sinensis* using ab initio-based, homology-based and transcriptome-assisted methods. The AUGUST and SNAP(Korf 2004) were implemented to predict genes in the de novo approach. The protein sequences of *Callorhinchus_milii*, *Danio_rerio*, *Latimeria_chalumnae*, *Lepisosteus_oculatus*, and *Petromyzon_marinus* (Ensembl-100 version) were downloaded from the Ensembl database for homology-based gene set prediction by Exonerate software. Gene structures were annotated using three approaches (ab initio predictions, homologue proteins, and transcriptome data) that were combined using MAKER software. Furthermore, gene functions were annotated against seven public databases including the NCBI non-redundant protein sequences (Nr), Swiss-Prot, Kyoto Encyclopedia of Genes and Genomes (KEGG), Cluster of Orthologous Groups of proteins (COG), TrEMBL, InterPro and GO databases according to the best match of the alignments using blastp (E-value < $1 \times 10^{-5}$).

**Genome collinearity analysis**

MCscan (Python version)(Tang, et al. 2008) was used for the genomic analysis between *A. sinensis*, *Atractosteus spatula*, *Acipenser ruthenus*, and *Psephurus gladius* with default parameters. The collinearity figure was drawn based on the gene collinear pair information between species by JCVI (https://github.com/tanghaibao/jcvi) or by Circos(Krzywinski, et al.).

**Gene family identification and phylogenetic tree construction**

To reduce the interference caused by whole genome duplications (WGD), we defined a criterion for screening gene families to construct a phylogenetic tree. The phylogenetic tree was constructed with 2,096 genes using PhyML(Guindon, et al. 2010).

A total of 13 species (*Acipenser sinesis*, *Acipenser ruthenus*, *Callorhinchus milii*, *Danio rerio*, *Gadus morhua*, *Latimeria chalumnae*, *Lepisosteus oculatus*, *Oryzias latipes*, *Petromyzon marinus*, *Polyodon spathula*, *Polypterus senegalus*, *Cyprinus carpio*, and *Salmo salar*) were applied in family clustering and the phylogenetic analyses. The protein-coding genes of the other 12 species were downloaded and filtered, and only the longest ORF with genes encoding more than 50 amino acids remained for the gene families clustering and phylogenetic analysis. Gene families were identified using Orthofinder(Emms and Kelly 2019). For the polyploidy species, *Polyodon spathula*, *Acipenser sinesis*, *Acipenser ruthenus, Cyprinus carpio*, and *Salmo salar*, we filtered the gene family with copy number setting to 1 or 2 of them. For *Cyprinus carpio* and *Salmo salar,* we randomly selected one copy as a single-copy orthologous gene. For *Polyodon spathula*, *Acipenser sinesis*, and *Acipenser ruthenus*, we selected a group (copy number of 1:1:1) with the least divergence as single-copy orthologous genes. Combining the other species, the single-copy orthologous genes (2,096) from gene families were further aligned using MUSCLE (MUSCLE, RRID: SCR_011812, version 3.8.31)(Edgar 2004) with default parameters and subsequently reverse-translated into codon sequences. These aligned sequences were concatenated to generate a super alignment matrix for phylogenetic reconstruction based on PhyML (PhyML, RRID: SCR_014629)(Guindon, et al. 2010) with proteins of the single-copy orthologs shared among the

13 species. Additionally, we employed IQ-TREE (v1.6.12) (Kalyaanamoorthy, et al. 2017) to construct gene phylogenetic trees of the single-copy orthologs and use Astral (v5.6.1) (https://github.com/maryamrabiee/Constrained-search) to integrate the gene trees. The resulting phylogenetic tree was consistent with the tree generated by PhyML.

**Karyotype analysis of *Acipenser sinensis***

Cultured muscle cells grown to 80-90% confluency cells were used for chromosome analysis. Briefly, cells were cultured in MEM medium with 20% FBS at 25 °C incubators and treated with colchicine solution (Sigma) at a final concentration of 20 μL/mL overnight. Successively, cells were trypsinized, centrifuged and gently resuspended in 0.75M KCl for 30-40 min at a 30 °C water bath. Then, cells were fixed with a 3:1 fixed solution of methanol and acetic acid three times for 15 min each. Finally, the resuspension of muscle cells was dropped onto a pre-cooled glass slide and stained with 10% Giemsa solution (Solarbio, China) for 10 min. The glass slides were calculated and 100 chromosome metaphase images were obtained under a light microscope (Leica, Germany).

**Screening of microsatellite markers**

SSRs were detected in unigenes by MIcroSAtellite (MISA; http://pgrc.ipk-gatersleben.de/misa, version 1.0) software. The following types of SSRs were detected: mono-, di-, tri-, tetra-, penta-, and hexa-nucleotide repeats, as well as compound SSRs [the sequence including more than one type of repeat units, e.g., (GA)n(TC)m]. Here, the standards for the repeat unit numbers were set as follows: mono-10, dimer-6, trimer-5, tetramer-5, pentamer-5, and hexamer-5. Tetramers were screened and used for primer selection. In this study, four-base repeat units were screened from the genome, and primers were synthesized and amplified to select primers that could amplify the target band to amplify target bands. The PCR products of 62 *A. sienensis* were performed in polyacrylamide gel electrophoresis to select the loci with polymorphism. The 5'-end of the upstream F primers of the polymorphic loci were labeled with FAM, HEX and TMRMA, respectively, and fluorescence typing was performed. Based on the results of the genotyping effect of these microsatellite loci (whether the peak pattern was single and clean, no stray peaks, multiple peaks, mismatched peaks, etc.), the loci with better typing effect were selected.

To select the loci that can be stably amplified in the *A. sinensis* genome, the locus stability test is necessary. In this study, 15 samples were randomly selected for three repeated experiments. Through repeated experiments and comparative analysis of genotyping results, microsatellite loci that were not stably amplified were eliminated. Following the genotyping peak map results, we checked whether there was variation in the allele length, and eliminated the data with a large difference in the length of the target sequence. The microsatellites used in this study were all four-base repeat sequences, so the allele size in the genotyping results should be different by 4 bp or an integer multiple of 4 bp. If not, the sequences will be eliminated. Finally, statistical analysis of the different methods was performed on the genotyping results of the 62 individual samples collected in this study. First, we used the MICRO-CHECKER software(Van Oosterhout, et al. 2004) to examine the results of the genotyping data to determine whether the locus had zero alleles and whether it conformed to a neutral mutation pattern, and eliminated those with a high zero allele frequency of microsatellite markers. Then, the polymorphism levels of these loci were assessed, and the Hardy-Weinberg equilibrium (HWE) and Linkage disequilibrium (LD) tests were performed to detect whether these microsatellite markers deviate from Hardy-Weinberg equilibrium and whether there is linkage disequilibrium between different loci. We screened 3,303 candidate sequences, and further 25 microsatellite markers were screened to further determine the ploidy of *A. sinensis*.

**Screening of the maximum number of alleles**

Ploidy of the species was estimated by the maximum number of alleles per individual at microsatellite loci. To obtain available simple sequence repeat (SSR) markers for determining ploidy of *A. sinensis*, all screened SSRs of a tetra-nucleotide repeat were verified by polyacrylamide gel electrophoresis (PAGE) and capillary electrophoresis on the ABI 3730 Genetic analyzer (ABI, US), respectively.

**Heterozygosity, allele frequency, and allele depth analysis**

To detect the heterozygous sequence polymorphism, paired-end reads of *A. sinensis* (~142× coverage) and *A. ruthenus* (~56× coverage, NCBI Accession: SRP174514) were mapped to their assembled scaffolds by aligner BWA (Version 0.7.12-r1039) and Samtools (Version 1.4). The heterozygous SNPs were called by Freebayes (v0.9.10-3-g47a713e) and filtered by following 4 thresholds: 1) the ratio of two alleles: *A. sinensis* (1:19-19:1), *A. ruthenus* (1:9-9:1); 2) the highest sequencing depth of SNP position: *A. sinensis* < 300×, *A. ruthenus* < 150×; 3) the lowest sequencing depth for each allele ≥ 6; and 4) the minimum distance for adjacent SNPs ≥ 5 bp. For each SNP site, there are three types of read depth (reference allele, alternative allele, and both). Then, we calculated both allele depth and the reference allele frequency (reference depth divided by both depths) of each SNP site. Finally, the density distribution of the allele depth and the allele frequency was counted and plotted (Fig. 2), where the peak in the largest depth of the distribution was defined as the depth of diplotype (4n for *A. ruthenus* 46X and 8n for *A. sinensis* 150X respectively). The SNP site allele frequency and ploidy (n) were evaluated by the following equation:

$$\text{SNP site allele frequency:} 1/n = \frac{depth\ of\ reference\ alleles}{depth\ of\ both\ alleles};$$

$$\text{SNP site of } A.\ ruthenus \text{ ploidy: } n = \frac{4(depth\ of\ both\ alleles)}{depth\ of\ diplotype} = \frac{4(depth\ of\ both\ alleles)}{46};$$

$$\text{SNP site of } A.\ sinensis \text{ ploidy: } n = \frac{8(depth\ of\ both\ alleles)}{depth\ of\ diplotype} = \frac{8(depth\ of\ both\ alleles)}{150}$$

**Heterozygous *k*-mer pairs analysis**

To disentangle the genomic ploidy properties of *C. carpio*, *T. arcticus*, *M. sativa*, *A. sinensis*, and *A. ruthenus*, we extracted the haplotype structures from heterozygous *k*-mer pairs using the Smudgeplots pipeline(Ranallo-Benavidez, et al. 2020). First, we produced a *k*-mer frequency file by KMC(Kokot, et al. 2017) with *k* = 21 from trimmed reads. Then, we searched for all heterozygous pairs of *k*-mers which differed at exactly one nucleotide through a systematic scan of all input k-mers. To avoid sequencing errors with genomic *k*-mers, we filtered the *k*-mers with a depth of less than 14, which was the depth of the first trough in the *k*-mer frequency curve. Finally, we performed the R script of the pipeline, plotted the smudge pots, and estimated ploidy using the coverage file of heterozygous *k*-mers. This tool performed gymnastics with the heterozygous *k*-mer pairs by comparing the sum of *k*-mer pair coverages (CovA + CovB) to their relative coverage (CovA / (CovA + CovB)).

**Distinctive repeats cluster in the homoeologous**

For a species of allopolyploid, some repeats may be expanded distinctively at each progenitor of the subgenome due to independent evolution before allotetraploid hybrids. The subgenome-specific repeats were used to distinguish subgenomes of allotetraploid *Xenopus laevis*(Session, et al. 2016) and *Carassius gibelio*(Wang, et al. 2022). Here, we tried to classify the subgenome of this fish using this strategy.

To avoid the influence of homoeologous exchange across two subgenomes after the WGD event, we chose homoeologous blocks with collinearity (> 10 Mb) instead of whole chromosomes (Table S16). The filtered homoeologous block pair count was 10 and the total length was 400 Mb. Then, we classified the transposable elements into clusters according to the target sequences in the Repbase or de novo consensus library. We analyzed each TE family content in each block with bp/1 Mb as a unit. The block_A/block_B content ratio in homoeologous pairs was calculated and log transformed according to ($\log_2$((Content.A+1)/(Content.B+1))). Finally, we picked out the 20 groups with the largest folds of differences in heatmaps using the R package.

**Topological structure analysis**

We screened the two-copy gene families in the genomes of *A. sinensis*, *A. ruthenus*, and *P. spathula*. Sequence divergences were calculated based on the six gene families of the three Acipenseriforme species, and we selected a set of gene families with minimum divergence values. A total of 1,438 gene families exist two copies in *A. sinensis* (S), *A. ruthenus* (R), and *P. spathula* (P), but exist only one copy of gene families in *L. oculatus* were used to construct gene trees and implied the topology of the species.

**LORe and AORe analysis**

Under LORe, speciation precedes rediploidization, allowing independent ohnologue divergence in sister lineages sharing an ancestral WGD event. A phylogenetic implication of LORe is a lack of 1:1 orthology between ohnologue pairs from different lineages, leading to the definition of the term 'tetralog' to describe a 2:2 homology relationship between ohnologues in sister lineages. Order differences in differentiation and doubling, as well as the influence of species characteristics, will lead to differential enrichment of LORe and 'Ancestral Ohnologue Resolution' (AORe)(Martin and Holland 2014; Robertson, et al. 2017). Nevertheless, previous studies ignored the coefficient of LORe and AORe, and the accuracy of the phylogenetic tree was affected. AORe would greatly improve the accuracy of differentiation time based on independent calculations.

Combined with gene family identification and genome collinearity analysis, we identified the potential ohnologues with 2:2:2:1 in *Acipenser sinesis*, *Acipenser ruthenus*, *Polyodon spathula*, and *Lepisosteus oculatus*. The protein sequences of ohnologues were aligned using MUSCLE (v3.8.425)(Edgar 2004) with the default parameters. These alignments were subsequently converted into CDS alignment by tracing the coding relationships using pal2nal.v14(Suyama, et al. 2006). Gblocks (v0.91b)(Talavera and Castresana 2007) was employed to conduct further checks (trim) on the CDS alignments with parameters "-t = c". The trimmed alignments with lengths less than 150 bp were filtered and then transmitted to IQ-Tree(Kalyaanamoorthy, et al. 2017) to infer the gene tree with settings: -alrt 1000 -bb 1000. Each trimmed gene was subjected to a gene tree analysis in the same manner. DensiTree(Bouckaert and Heled 2014) was used to visualize the topologies of these trees.

**Ks analysis of AORe**

We calculated the Ks of AORe ohnologues for inter-species and intra-species in the three genomes. Ks analyses were performed using default parameters and the 'fasttree' node-weighting method in the wgd package(Edgar 2004; Yang 2007a; Guindon, et al. 2010; Zwaenepoel and Van de Peer 2019). Distributions in Ks were plotted based on node-averaged values as calculated in the wgd package. The Gaussian mixture models (GMMs) were fitted to the Ks distribution following the wgd pipeline, with the optimal number of components assessed using the Bayesian information criterion.

**Estimation of divergence time based on AORe**

The protein sequences of AORe in Acipenseriformes and other species ohnologues were extracted to estimate the divergence times between the species and the subgenome-like (ohnologue divergence after WGD) of Acipenseriformes using MCMCTree (Yang 2007b) by using the proteins with the approximate likelihood calculation method, the 'correlated molecular clock' and 'REV' substitution model, successively. Four data sets from the TimeTree database(Hedges, et al. 2006) were used for calibration.

**Unitary pseudogenes analysis**

Responding to polyploidization, either immediately (for neoallopolyploids) or after some "diploidization" period (for autopolyploids), disomy is re-established. Once this state is reached, polyploidy genomes are shaped by the longer-term evolutionary forces of mutation, drift, pseudogenization, selection, and recombination, leading to further differentiation of the homologous(Wolfe 2001; Session, et al. 2016).

Unitary pseudogenes are nonfunctional genes that decayed in their original place, whereas retroposed pseudogenes have been transposed to a new location(Zhang, et al. 2010; Session, et al. 2016). We proposed that some missing homoeologues would be present in the genome as unitary pseudogenes after the Ass4R event. To search for pseudogenes, we looked for gene families with quadrivalent pairing collinearity in *A. ruthenus* and *A. sinensis*. Each gene family conformed to the AORe model with two copies in *A. ruthenus* and one copy and one pseudogene in *A. sin*ensis. We first aligned the *A. ruthenus* genes to the *A. sinensis* genome using BLASTP with the parameters "-e 10-5". The best hit region of each gene with 1 Kb flanking sequence was cut down and re-aligned with the corresponding orthologous protein sequence using GeneWise(Birney, et al. 2004; Blanchette, et al. 2004) with parameters "-genesf -tfor -quiet", which could help to define the detailed exon-intron structure of each gene(Zhang, et al. 2010). All of the genes containing frame shifts or premature stop codons as reported by GeneWise were considered candidate pseudogenes. We further carried out a series of filtering processes: 1) Candidate pseudogenes that did not support collinearity to *A. ruthenus* were filtered, 2) Frame shift or premature stop codon sites in candidate pseudogenes with a low number of reads covering (< 5X) were considered as assembly errors and filtered, and 3) Mutated sites in candidate pseudogenes near exon-junction boundaries (< 6 bp) were filtered. Finally, the screened quadruple collinearity genes (2:2 in *A. sinensis* and *A. ruthenus*) with one pseudogene in the genomes of *A. sinensis* which conformed to the AORe model were adopted. In total, we identified 344 unitary pseudogenes in the genome.

To estimate the age of pseudogenes, we used an approach(Zhang, et al. 2010; Session, et al. 2016) that depends on the excess non-synonymous substitution of a pseudogene relative to normal functional orthologue with known divergence time. For each pseudogene in the quadruple collinearity genes, we filtered the mutated site to be translatable and then calculated the Ka and Ks with the AORe homologous gene. Another two coding homologous genes in the quadruple collinearity genes were used to calculate the Ka and Ks meanwhile. If the nonsynonymous substitutions Ka in the pseudogene lineage evolved as the same Ka/Ks ratio measured between the coding functional genes (another two coding homologous genes in the quadruple collinearity genes) up until the pseudogenization time T * (KsT) and at the synonymous rate Ks afterwards reflecting the relaxation of selection, we can express the pseudogenization time, in units of Ks, for each pseudogene.

$$KsT = (KaP - KaC/KsC \times KsP) / (1 - KaC/KsC)$$

KaP and KsP values were Ka and Ks of pseudogene and KaC and KsC values were Ka and Ks of coding functional genes.

### Figures S1 to S12

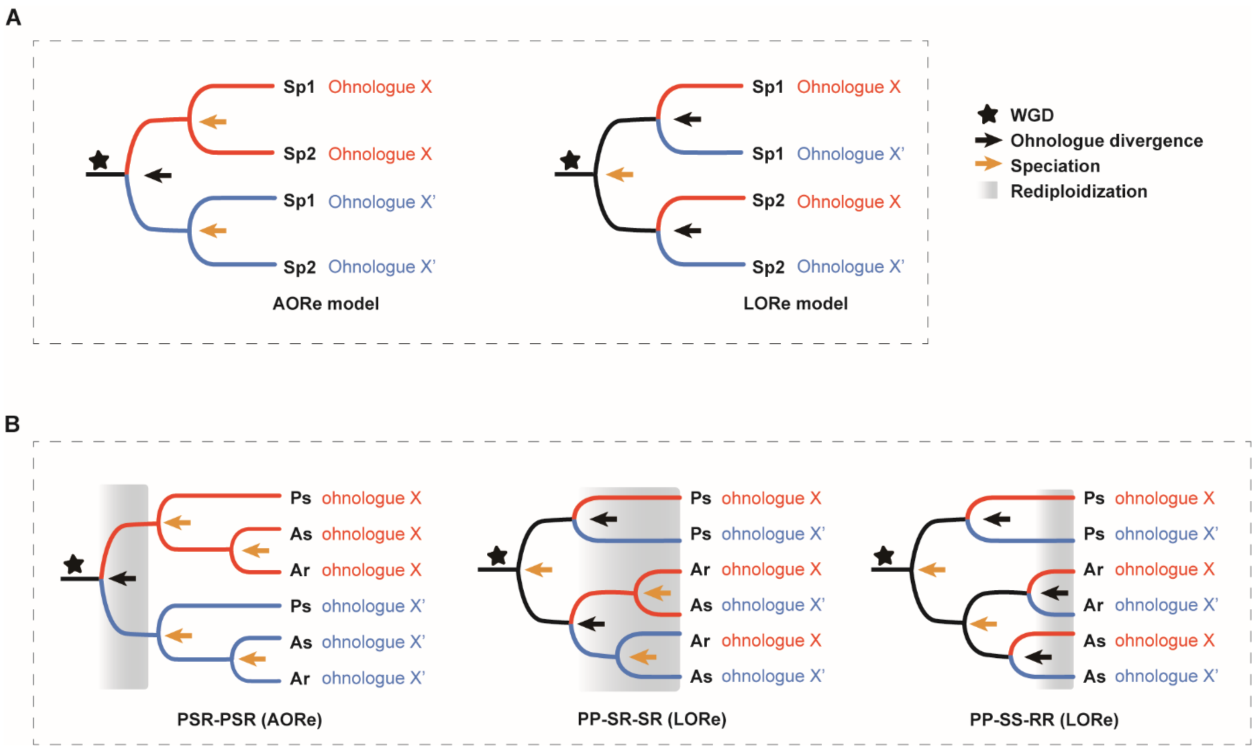


**Figure S1 Relationships among WGD, divergence, and rediploidization based on AORe and LORe model. (A)** The AORe model of post-WGD, and the LORe model of post-WGD evolution following delayed rediploidization. **(B)** PSR-PSR, PP-SR-SR, and PP-SS-RR model. P, S, and R represent Polyodon spathula, Acipenser sinensis, and Acipenser ruthenus, respectively. PSR-PSR was constructed by AORe and the divergence and speciation occurred after Acipenseriforme-specific WGD (As3R) and the complete rediploidization. PP-SR-SR was constructed by LORe and the two families of Acipenseriformes diverged after As3R before the complete rediploidization. PP-SS-RR was constructed by LORe and the divergence and speciation occurred before the complete rediploidization after As3R.

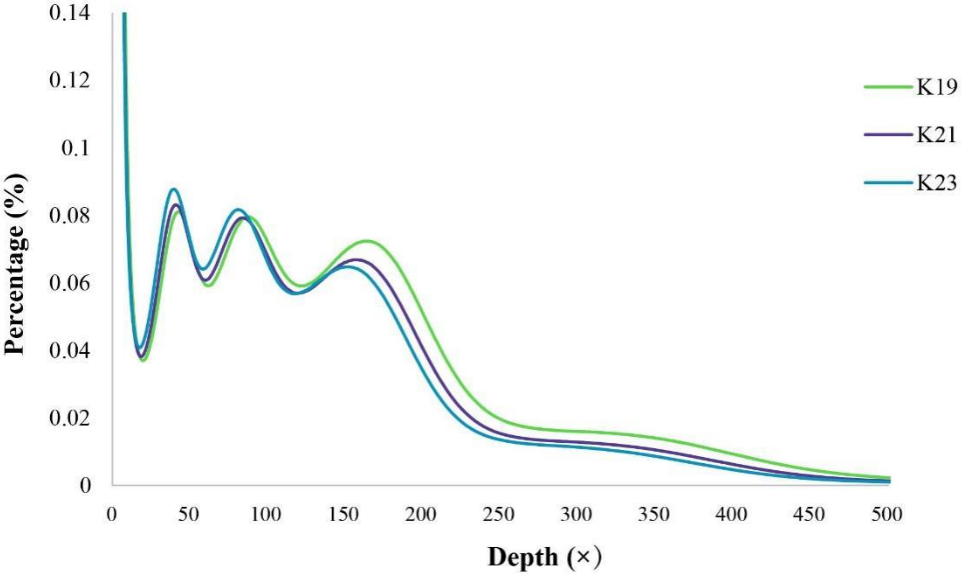


**Figure S2 19-mer, 21-mer, and 23-mer distributions were used for the estimation of genome size.** The X-axis represents the depth of sequencing. The Y-axis is the proportion of 17-mer frequencies at different depths. The total genome size was estimated according to the following formula: genome size = *k*-mer number/peak depth, where *k*-mer number is the total number of *k*-mers, and peak depth is the maximal frequency.

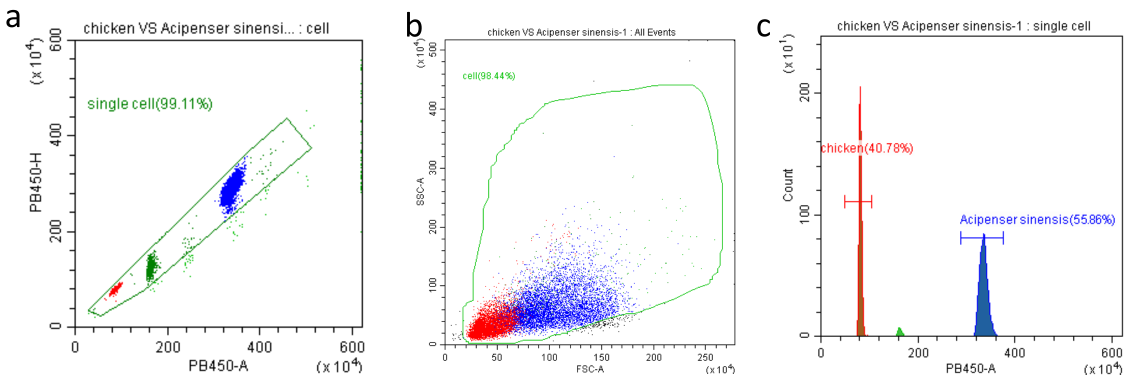


d

Tube Name: chicken VS Acipenser sinensis-1

Sample ID:

| Population | Events | % Total | % Parent | CV PB450-A | Median PB450-A | Mean PB450-A |
|---|---|---|---|---|---|---|
| All Events | 12880 | 100.00% | 100.00% | 58.67% | 3242204.3 | 2296296.5 |
| cell | 12679 | 98.44% | 98.44% | 55.50% | 3239217.8 | 2278178.0 |
| single cell | 12566 | 97.56% | 99.11% | 54.80% | 3237777.5 | 2257395.5 |
| chicken | 5124 | 39.78% | 40.78% | 3.23% | 809030.2 | 812057.1 |
| Acipenser sinensis | 7019 | 54.50% | 55.86% | 2.53% | 3335678.5 | 3339971.0 |

**Figure S3 Flow Cytometer result of *Acipenser sinensis*.**

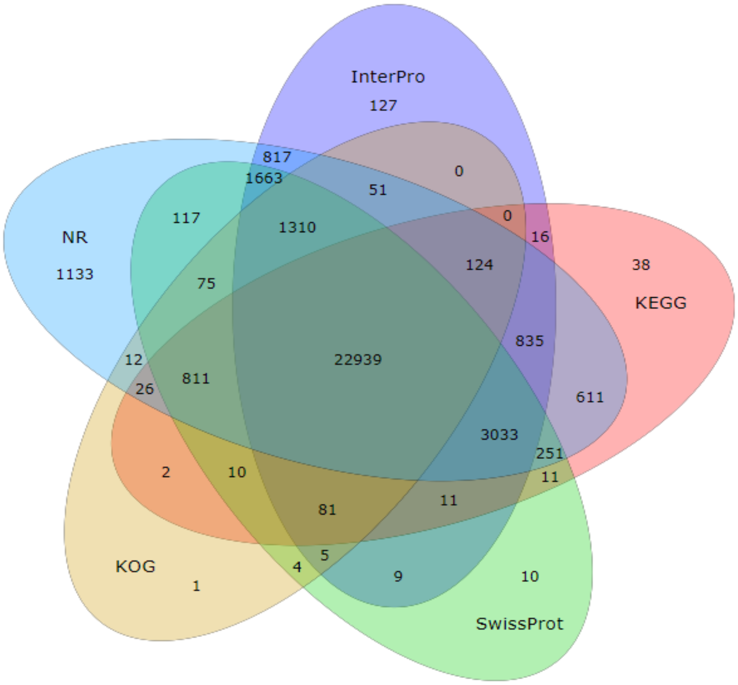


**Figure S4 Veen diagram of gene function annotation based on Nr, InterPro, KEGG, SwissProt, and KOG databases.** A total of 22,939 genes were shared by the eight database annotations.

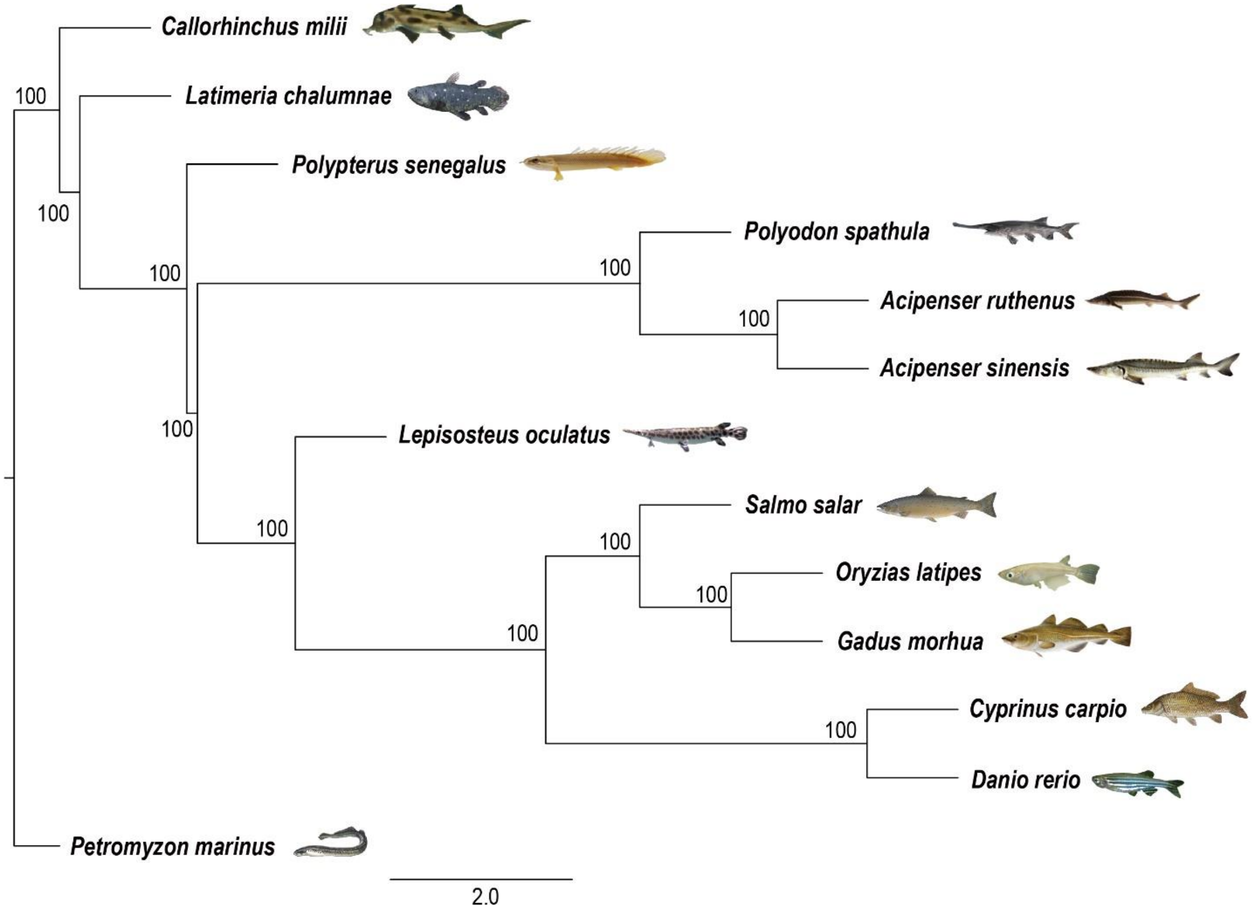


**Figure S5 Phylogenetic tree constructed with 2,096 single copy orthologous genes using Astral.**

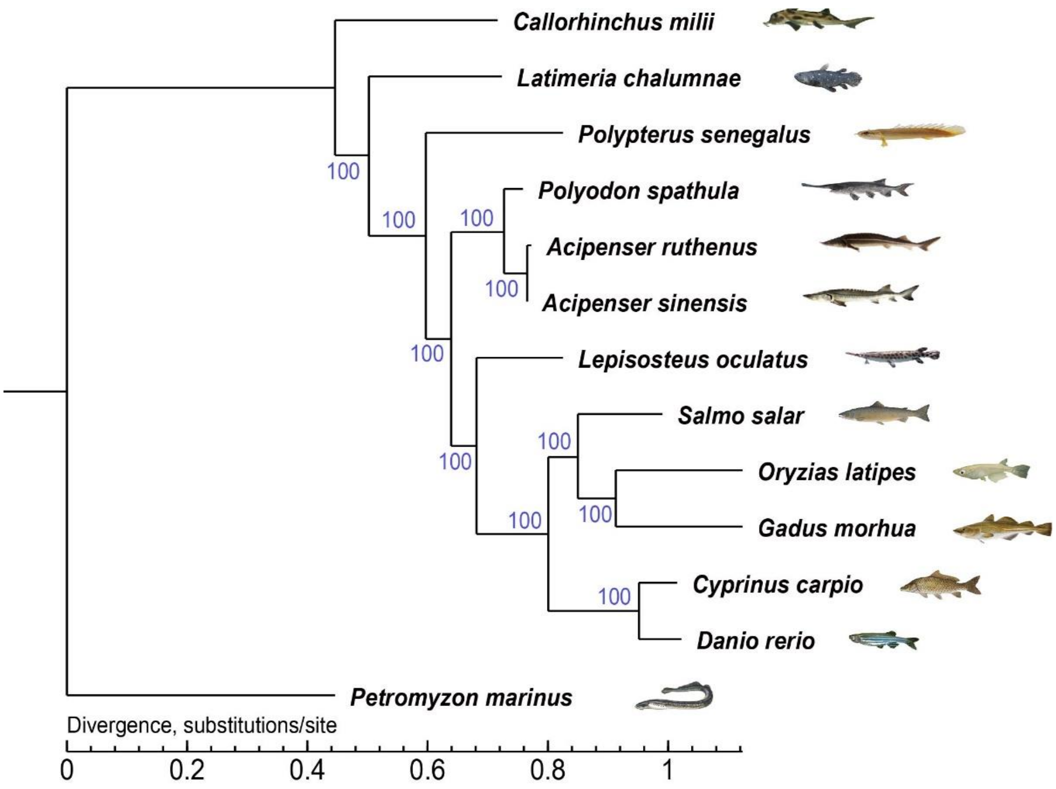


**Figure S6 Phylogenetic tree constructed with 2,096 single copy orthologous genes using PhyML.** The branch length represents the neutral divergence rate.

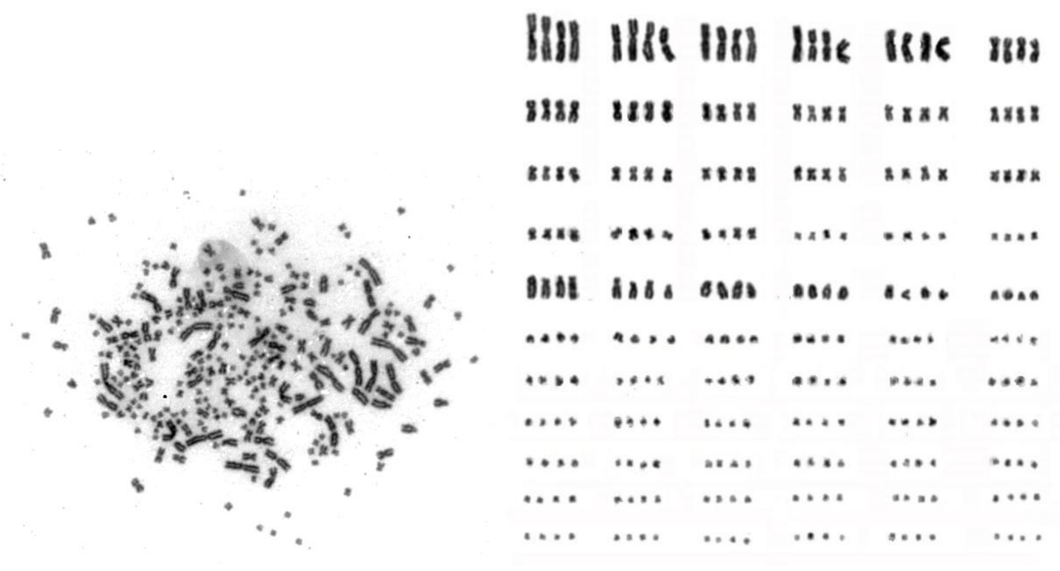

**Figure S7 Karyotype of *Acipenser sinensis*.** A total of 264 chromosomes were identified (left) and ranged in groups containing four chromosomes (right). The first row is the macro-chromosomes.

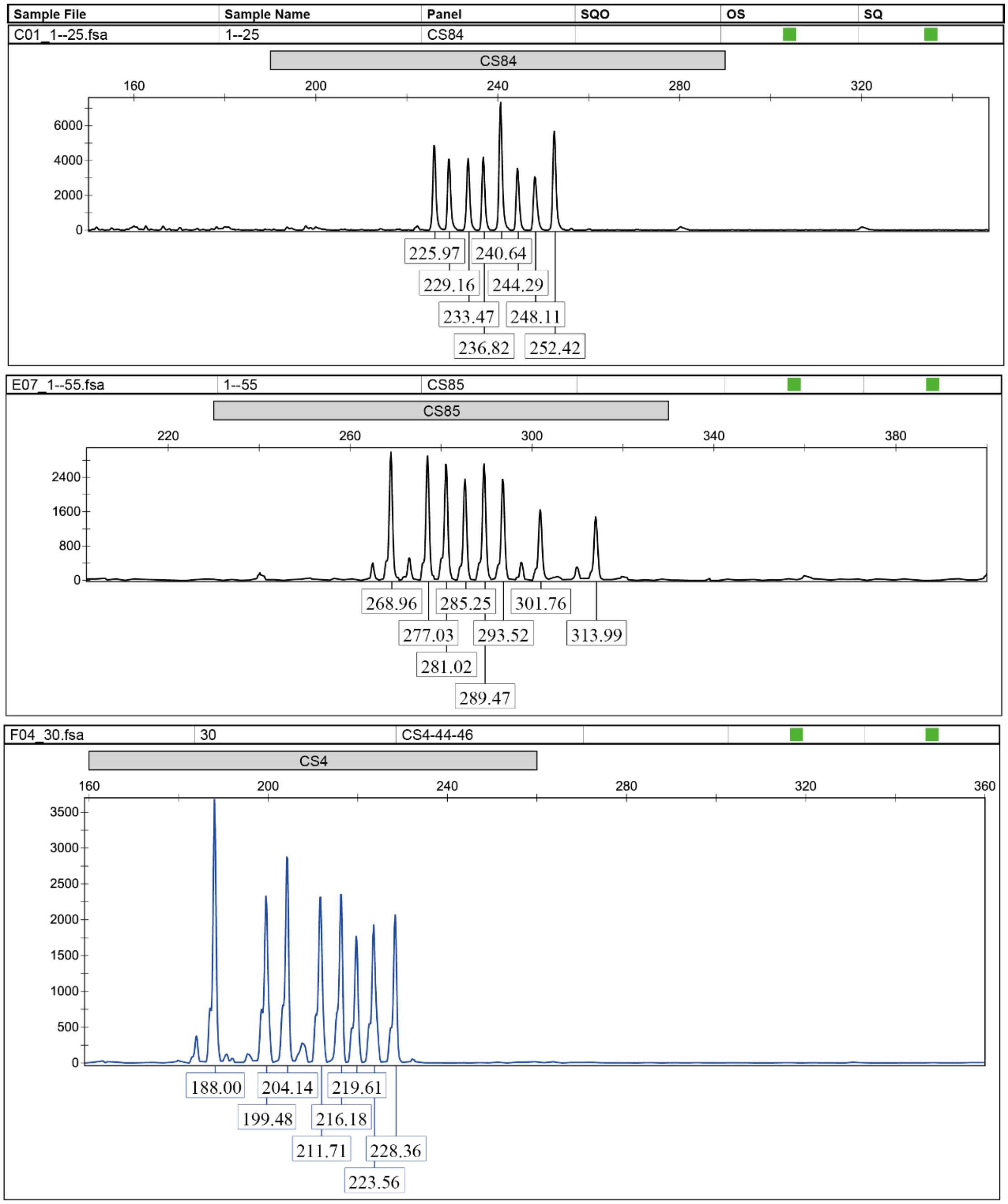


**Figure S8 Ploidy identification based on SSRs.** Eight peaks were identified by tetra-nucleotide repeat SSRs using capillary electrophoresis on ABI PRISM 3730 Genetic Analyzer.

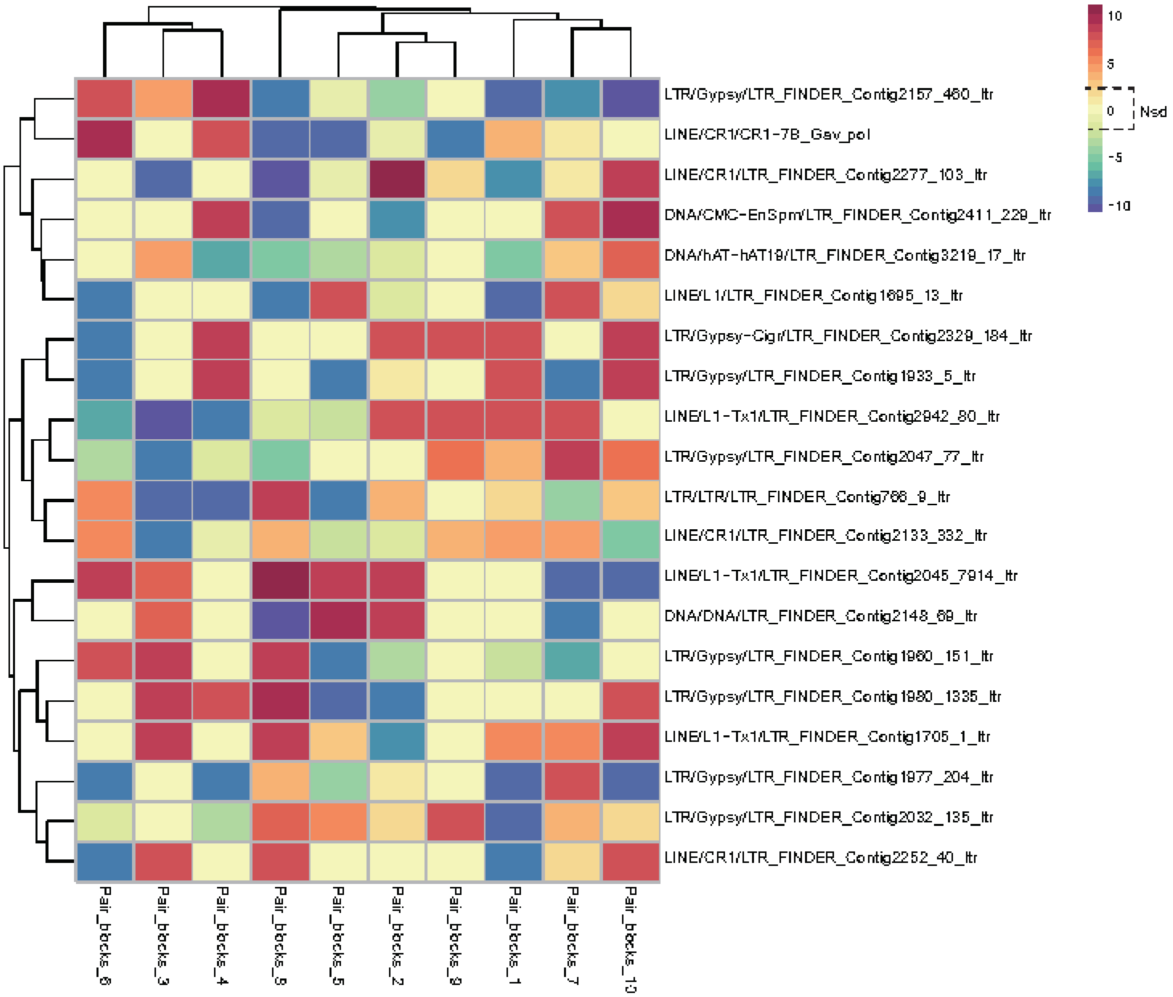


**Figure S9 Detection of homology and heterology based on differential TEs screening.** Pair_blocks_1-10 at the X-axis are 10 homoeologous sequence blocks with collinearity. The Y-axis represents 20 TEs in the 10 homoeologous blocks. Nsd represents that the block pair of TE has no significant difference. If at least one Nsd block appears in a row, the row is not a differential TE.

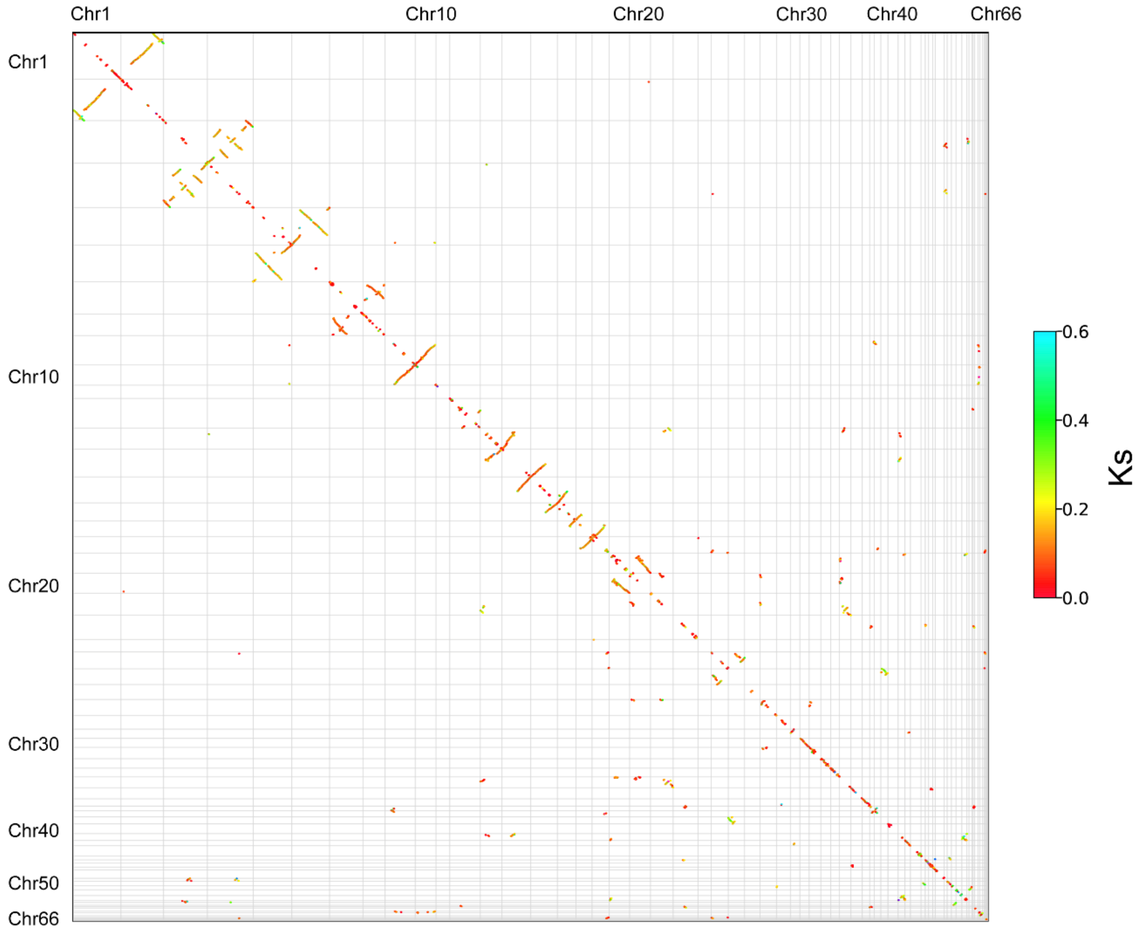


**Figure S10 Homologous gene dot plot within the *Acipenser sinensis* genome.** Ks value for homologous genes in each inferred collinear block is shown. The high Ks values were presented on macrochromosomes 1-6.

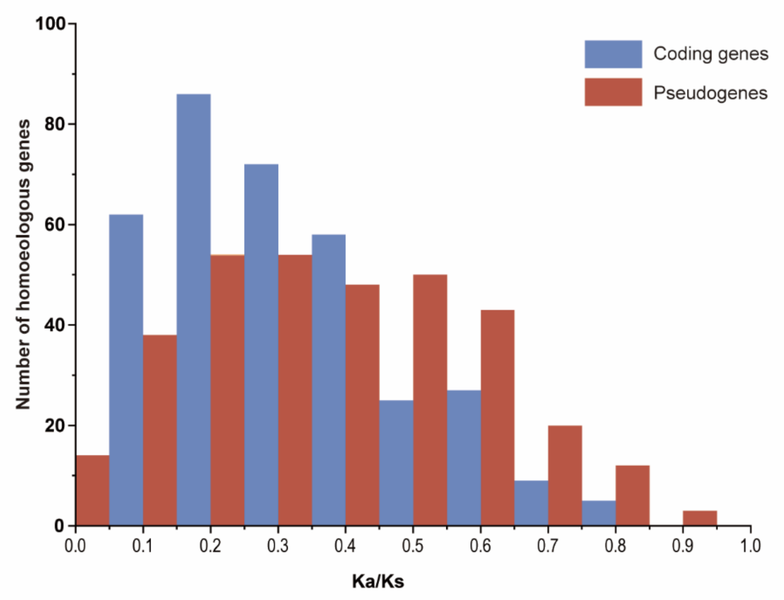


**Figure S11 Distribution of Ks analysis in coding genes and unitary pseudogenes.**

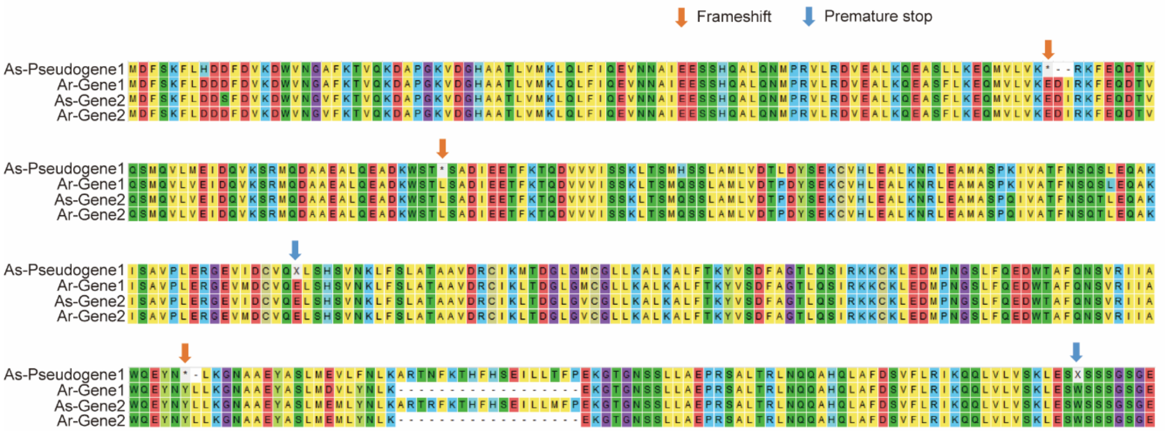


**Figure S12 Amino acid sequence of the COG7 coding gene and pseudogene alignment in *Acipenser sinensis* and *Acipenser ruthenus*.** Four genes were selected, As-Pseudogene1 (Gene id: ACSI009096-D2) and Ar-Gene1 (ACSI009096), As-Gene2 (XP_033886174.2) and Ar-Gene2 (XP_033907760.1) were from the common AORe, respectively. Missing codons are marked with dashes. Frameshifts and premature stops are marked by * and X, respectively (pointed by the red arrow and blue arrow).

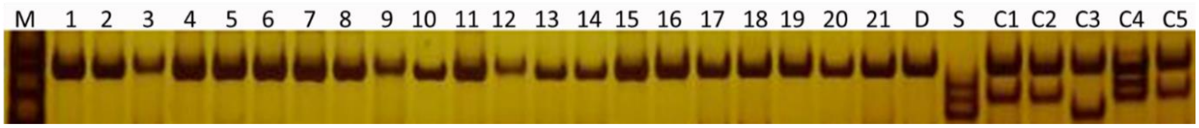


**Figure S13 Identification results for gynogenetic *Acipenser sinensis* by microsatellite DNA analysis on PAGE.** Lane M represents the DNA ladder marker. Lanes 1-21 show gynogenetic individuals. Lanes D and S represent the dam (maternal) and sire (paternal), respectively. Lanes C1-C5 represent the control diploid individuals.
